# VPS13C/PARK23 initiates lipid transfer and membrane remodeling for efficient lysosomal repair

**DOI:** 10.1101/2025.10.23.684214

**Authors:** Oluwatobi Andrew Adeosun, Christian Schröer, Elisabeth Südhoff, Fabio Bergenthal, Katharina Sommer, Emely Döffinger, Britta Fiedler, Angelika Hilderink, Ann-Katrin Lehmann, Lea-Sophie Pohle, Florian Fröhlich, Kenji Maeda, Rainer Kurre, Michael Holtmannspötter, Bianca M. Esch, Joost C. M. Holthuis

## Abstract

Perturbations in lysosome integrity are tightly linked to neurological disorders and ageing, but the underlying pathogenic mechanisms are incompletely understood. Using an unbiased proteomic approach, we here identified the bridge-like lipid transport protein VPS13C/PARK23 as a key component of a global early response pathway to lysosome damage. VPS13C readily binds lysosomes under mechanical or osmotic tension in anticipation of membrane lesions. The latter trigger a conformational change in the protein’s *C*-terminus, involving its ATG2C domain acting as sensor of damage-induced lipid packing defects. We show that ER-lysosome contacts formed by VPS13C provide critical binding platforms for OSBP/ORPs to enable efficient ER wrapping of damaged lysosomes. A chemical approach to assess directional ER-to-lysosome lipid transport revealed that VPS13C is essential for large-scale lipid delivery to acutely damaged lysosomes to facilitate their repair. Our findings offer new mechanistic insights into how loss-of-function mutations in *VPS13C* may enhance the risk of Parkinson’s disease.

## INTRODUCTION

Lysosomes are essential degradative organelles with central roles in nutrient sensing, cell growth signaling, cellular stress handling, and immunity^1^. For their catabolic function, lysosomes rely on dozens of hydrolytic enzymes in their acidified lumen, which is bound by a single limiting membrane that is lined on the luminal side with a protective glycocalyx^2^. A key vulnerability of lysosomes is the damage inflicted by diverse materials such as incoming pathogens that seek access to the cytoplasm, endocytosed silica crystals that can puncture the membrane, or lysosomotropic substances that intercalate into or otherwise destabilize the lipid bilayer^3,4^. Damage to the lysosome-limiting membrane can also be caused by endogenous conditions, such as lipid peroxidation or the accumulation of lipid metabolites in lysosomal storage disorders^5–7^. Perturbations in lysosome integrity are frequently associated with neurodegenerative disorders and can have deleterious consequences, as the release of hydrolytic enzymes into the cytosol can induce inflammatory signaling, autophagy dysfunction and cell death^6,8^.

To maintain lysosome integrity, cells are equipped with multiple lysosome damage response pathways that are activated pending on the degree of injury. When damage is severe, involving membrane rupture, lysosomes are marked for degradation by lysophagy. This process is initiated by the recruitment of cytosolic galectins and glycoprotein-specific ubiquitin ligases to abnormally exposed luminal glycans, resulting in engulfment of the damaged lysosome by autophagic membranes^9,10^. Lysophagy is a slow process in which the whole organelle is ultimately sacrificed^11^. At the same time, an mTORC1-governed signaling pathway senses the damage and induces gene expression for lysosomal components to enable the biogenesis of new lysosomes^4,12,13^. While removal and replacement of damaged lysosomes are essential for maintaining cellular homeostasis, these pathways fall short in efficiently neutralizing the harmful effects of acute lysosomal leakage. For this purpose, cells evolved diverse repair mechanisms to instantly clear minor lesions from the lysosome-limiting membrane, preventing mildly injured lysosomes from rupturing and being sacrificed.

One lysosomal repair mechanism is mediated by the endosomal sorting complexes required for transport (ESCRT) machinery^14^. ESCRT proteins are organized in functionally distinct complexes that assemble in response to Ca^2+^ efflux at sites of membrane perforation. Concentric polymerization of ESCRT-III proteins subsequently drives an invagination of the lesion sites and, with help of the AAA ATPase VPS4, sorts them into intraluminal vesicles for degradation^14–16^. Clearance of lesions from the lysosome-limiting membrane can also be achieved via a sphingolipid-dependent repair pathway. Various conditions perturbing lysosome integrity trigger a rapid Ca^2+^-activated scrambling and cytosolic exposure of sphingomyelin (SM)^17,18^. On the cytosolic surface, SM is then cleaved by neutral sphingomyelinase to ceramide, a cone-shaped lipid that occupies a smaller membrane area than SM. Ceramides released by SM turnover readily cluster in microdomains that have a negative spontaneous curvature^19,20^, which would promote an inverse budding of the damaged lysosomal membrane region in a process analogous to ESCRT-mediated formation of intraluminal vesicles^18^.

For effective removal of membrane lesions via sphingolipid- and ESCRT-mediated repair pathways, injured lysosomes would require a constant supply of fresh lipids to sustain intraluminal vesicle formation. Lysosome damage triggers the formation of ER-lysosome contact sites mediated by ER-anchored OSBP/ORP family lipid transport proteins in response to a damage-induced production of phosphatidylinositol-4-phosphate (PI4P) on the lysosomal surface via the phosphoinositide-initiated membrane tethering and lipid transport (PITT) pathway^21^. As OSBP and ORPs selectively transport cholesterol and phosphatidylserine (PS) from the ER to the lysosome-limiting membrane in exchange for PI4P, they do not mediate net lipid transfer to damaged lysosomes. However, the rise in lysosomal PS levels stimulates the recruitment of the bridge-like lipid transport protein ATG2^21^. Unlike OSBP and ORPs, which selectively bind one lipid at a time for transport, ATG2 has a hydrophobic groove along which multiple lipids can slide simultaneously to facilitate bulk lipid transfer from the ER to lysosomes^22^. This large-scale lipid transport by ATG2 is initiated relatively late in the cellular response to lysosome damage as it relies on the gradual accumulation of PS. Wang *et al.*^23^ recently reported that VPS13C, another bridge-like lipid transport protein encoded by the Parkinson’s disease gene *PARK23*, is acutely mobilized to damaged lysosomes where it tethers their membranes to the ER. VPS13C recruitment does not require PITT pathway activation or Ca^2+^ efflux but instead relies on a signal originating from the damaged lysosome surface to release an auto-inhibited conformation of the protein that prevents its access to lysosome-bound Rab7. Loss of VPS13C enhances lysosome fragility, suggesting a role for the protein in lipid delivery to support lysosome repair^23^. However, this prediction awaits experimental validation. Another major outstanding question is whether VPS13C- and OSBP/ORP/ATG2-dependent lipid transport pathways function independently of each other or are hierarchically organized to maintain lysosomal integrity.

Using an unbiased proteomic approach, we here identified VPS13C as a core component of an early protective cellular response to lysosomal damage. We show that VPS13C readily binds mechanically or osmotically stressed lysosomes prior to the manifestation of membrane damage. In response to the latter, lysosome binding of VPS13C becomes strictly dependent on the amphiphilic properties of its *C*-terminal ATG2C domain, which mediates recognition of damage-induced lipid packing defects. Loss of VPS13C abolished both recruitment of OSBP/ORP proteins and ER wrapping of damaged lysosomes. Using a chemical strategy for imaging intracellular lipid flows in real time, we find that VPS13C is essential for initiating large-scale ER-to-lysosome lipid transfer to facilitate efficient repair of acutely damaged lysosomes. Our data collectively indicate that VPS13C acts upstream of OSBP/ORP/ATG2 to coordinate lipid transport in synchrony with lysosomal damage and repair, hence providing a mechanistic basis for how loss-of-function mutations in *VPS13C* may disrupt lysosomal integrity and contribute to the development of Parkinson disease.

## RESULTS

### A proteome-wide screen for lysosomal damage markers yields VPS13C/PARK23

To further elucidate the cellular mechanisms that mediate efficient lysosomal repair, we conducted an unbiased proteome-wide search for proteins that are rapidly recruited to damaged lysosomes. Lysosomes were affinity-purified from HeLa cells expressing GFP-tagged LAMP1 using paramagnetic beads decorated with an anti-GFP antibody (**Fig. 1a, b**). Lysosomal damage was induced by treating cells with the lysosomotropic reagent l-Leucyl-l-leucine methyl ester (LLOME; 500 μM, 30 min), which is membrane permeable and rapidly condenses in lysosomes to membranolytic poly-leucine that perforates the limiting membrane^24^. The purity of lysosome isolates from control and LLOMe-treated cells was assessed by immunoblotting, which showed that both isolates contained readily detectable levels of LAMP1 but were largely devoid of protein markers of the ER (calnexin), PM (Na^+^/K^+^-ATPase), mitochondria (pMito60) and cytosol (β-actin) (**Fig. 1c**). To determine the proteomic landscapes of intact and damaged lysosomes, we used high-resolution data-independent acquisition (DIA) liquid chromatography/tandem mass spectrometry (LC/MS-MS). A comparative proteomic analysis of lysosome isolates from LAMP1-GFP and background cells confirmed an efficient purification of lysosomes with minimal contamination from other organelles, irrespective of LLOMe treatment (**Suppl. Fig. 1a, b**). Isolates from untreated cells displayed a highly significant enrichment of both integral membrane and luminal lysosomal proteins, suggesting the purification of intact lysosomes. However, isolates from LLOMe-treated cells were relatively depleted in luminal lysosomal proteins (**Fig. 1d; Suppl. Fig. 1c**), presumably because perforation of the lysosome-limiting membrane caused their partial release.

**Figure 1.**
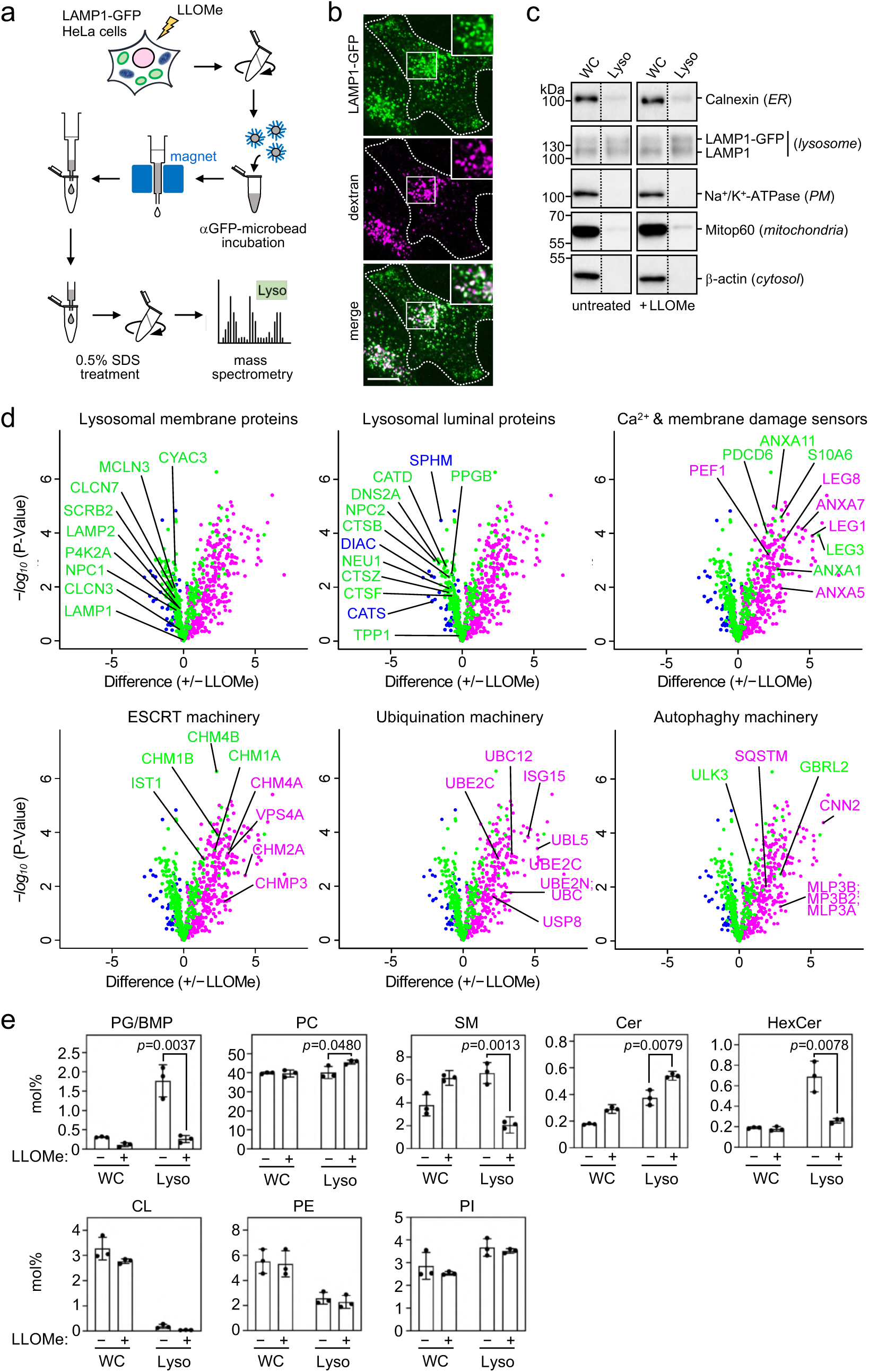
Damage-induced changes in lysosomal protein and lipid profiles. (**a**) Workflow for affinity purification of lysosomes from untreated and LLOMe-treated (500 μM, 30 min) HeLa cells stably transduced with GFP-tagged LAMP1. (**b**) HeLa cells expressing LAMP1-GFP (*green*) and fed Alexa Fluor647-labelled 10kDa dextran (*magenta*) for 16 h were imaged by spinning disk microscopy. Scale bar, 10 µm. (**c**) Whole cell lysates (WC) and lysosomes purified from cells treated as in (a) were subjected to immunoblot analysis using antibodies against calnexin (ER), LAMP1 (lysosomes), Na+/K^+^-ATPase (PM), mitop60 (mitochondria), and β-actin (cytosol). Samples were loaded on an equivalent basis. (**d**) Volcano plots showing enrichment and depletion of selected proteins on damaged lysosomes grouped by their molecular function. Fold changes were calculated from three independent biological replicates and plotted on the x-axis against the negative logarithmic *P*-values on the y-axis. *Green*, proteins enriched in both control and damaged lysosomes; *blue*, proteins selectively depleted in damaged lysosomes; *magenta*, proteins selectively enriched in damaged lysosomes. (**e**) Lipid composition of whole cell lysates (WC) and lysosomes purified from cells treated as in (a) was determined by mass spectrometry-based shotgun lipidomics. Levels of the different lipid classes are expressed as mol% of total identified lipids. BMP/PG, (bis(monoacyl-glycerol)phosphate/phosphatidylglycerol; PC, phosphatidylcholine; SM, sphingomyelin; Cer, ceramide; HexCer, hexosylceramide; CL, cardiolipin; PE, phosphatidylethanolamine; PI, phosphatidylinositol.

Consistent with an efficient purification of lysosomes, shotgun lipidomics revealed that lysosome isolates from untreated cells were devoid of the mitochondrial lipid marker cardiolipin (CL) and enriched in bis(monoacyl-glycerol)phosphate (BMP), a lipid primarily found in lysosomal intraluminal vesicles and here quantified together with the isobaric phosphatidylglycerol (PG) and reported as BMP/PG (**Fig. 1e**). Notably, LLOMe treatment caused a dramatic decrease in BMP/PG levels, possibly due to a damage-induced rupture of lysosomes and subsequent loss of intraluminal vesicles. LLOMe treatment also caused a significant rise in lysosomal ceramide levels and concomitant drop in SM and hexosylceramide (HexCer) levels, with the lipid species affected sharing the same acyl chain composition (**Suppl. Fig. 1d**). These results are largely in agreement with our previous finding that lysosomal damage triggers SM scrambling and hydrolysis to promote lysosomal repair by a ceramide-dependent formation of intraluminal vesicles^18^.

Proteins selectively enriched in LLOMe-damaged lysosomes were classified into different functional groups (**Figs. 1d, 2a**). The group of Ca^2+^ and membrane damage sensors included several annexins (ANXA1-7, 11), a family of Ca^2+^-regulated phospholipid-binding proteins implicated in sealing of membrane lesions^25^, and galectins (LEG1, 3, 7, 8), sugar-binding lectins that recognize abnormally exposed luminal glycans at sites of membrane rupture and help orchestrate membrane repair or initiate autophagy to clear membrane remnants^26^. Another prominent group comprised components of the ESCRT machinery and included ESCRT-III subunits (CHMP1-4 and IST1) and the ATPase VPS4A, which cooperate to clear lesions in the lysosome-limiting membrane via an inverse budding and fission process^14,15^. Additionally, we identified numerous components of the ubiquitin and autophagy machineries, many of which are involved in marking damaged lysosomes for lysophagy to mediate their destruction^4^.

**Figure 2.**
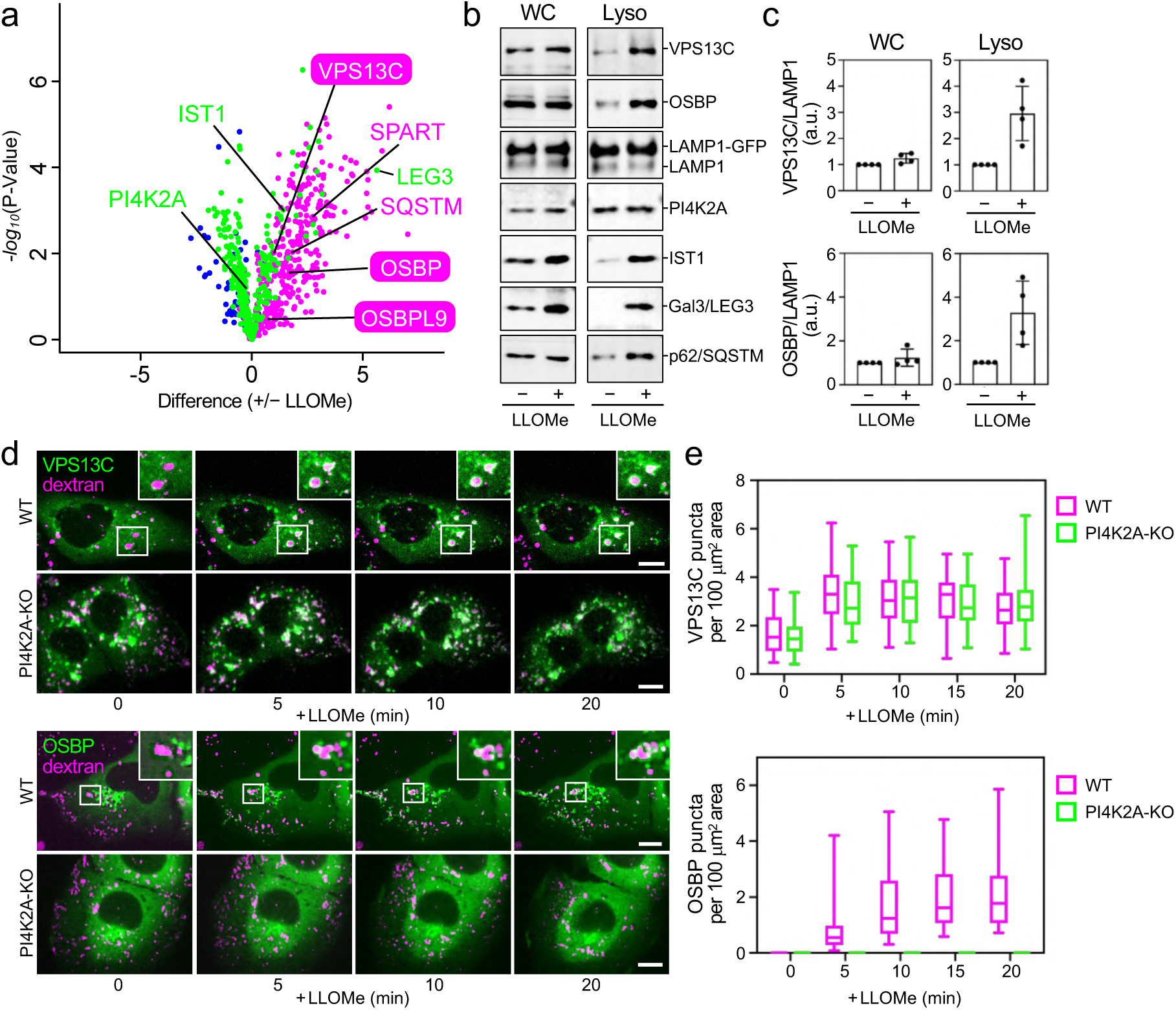
Recruitment of VPS13C and OSBP to damaged lysosomes occurs via distinct mechanisms. (**a**) Volcano plot showing enrichment of VPS13C, OSBP and various lysosomal damage markers on lysosomes affinity-purified from HeLa cells treated with LLOMe (500 μM, 30 min). *Green*, proteins enriched in both intact (–LLOMe) and damaged (+LLOMe) lysosomes; *blue*, proteins selectively enriched in intact lysosomes; *magenta*, proteins selectively enriched in damaged lysosomes. (**b**) Whole cell lysates (WC) and lysosomes purified from untreated and LLOMe-treated HeLa cells (500 μM, 30 min) were subjected to immunoblot analysis using antibodies against VPS13C, OSBP, LAMP1, PI4K2A, hIST1, Gal3/LEG3 and p62/SQSTM. (**c**) Quantification of the data in (b). Bars show normalized values relative to control. Data are means ± SD. *n* = 4 biological replicates. (**d**) Time-lapse fluorescence images of U2OS wildtype (WT) and PI4K2A-KO cells expressing VPS13C-mClover or OSBP-GFP (*green*) that were fed CF633-labelled 70kDa dextran (*magenta*) and treated with 1 mM LLOMe for the indicated time. Cells were imaged by spinning disk microscopy. Scale bar, 5 µm. (**e**) Time-course plotting VPS13C- and OSBP-positive puncta per 100 μm^2^ area in cells treated as in (d). Data are means ± SD (VPS13C: *n* = 31 cells for WT, 29 cells for PI4K2A-KO; OSBP: *n* = 35 cells for WT, 33 cells for PI4K2A-KO) from 2 independent experiments.

We also identified multiple components of the PITT pathway (PI4K2A, OSBP, OSBPL9/ORP9) as well as the bridge-like lipid transport protein VPS13C/PARK23 (**Fig. 2a**). The accumulation of OSBP and VPS13C on LLOMe-damaged lysosomes was validated by immunoblot analysis of lysosome isolates (**Fig. 2b, c**) and by confocal life-cell imaging of U2OS cells expressing GFP-tagged OSBP and mClover-tagged VPS13C (**Fig. 2d**). Under resting conditions, OSBP-GFP was exclusively localized in the Golgi complex region, while an additional pool was diffusely distributed throughout the cytosol. Upon addition of LLOMe, OSBP gradually accumulated on CF633-dextran-positive organelles, reaching a plateau at 10-15 min of LLOMe treatment (**Fig. 2e**). OSBP recruitment to CF633-dextran-positive organelles was completely abolished in PI4K2A-KO cells (**Fig. 2d, e; Suppl. Fig. 2a**), consistent with previous studies showing that lysosomal translocation of OSBP requires PI4P generated by PI4K2A on the surface of damaged lysosomes^21,27^. Confocal imaging of VPS13C-mClover expressing U2OS cells under resting conditions showed a few puncta or doughnut-like structures that often co-localized with CF633-dextran-positive organelles while the bulk of the protein displayed a diffuse cytosolic distribution. LLOMe addition caused a rapid recruitment of cytosolic VPS13C to CF633-dextran-positive organelles, which reached a plateau at 5 min of LLOMe treatment (**Fig. 2d, e**). VPS13C was still efficiently recruited to CF633-dextran-positive organelles in PI4K2A-KO cells in response to LLOMe, even though these cells appeared to mobilize a larger fraction of the protein to lysosomes under resting conditions. The observation that recruitment of VPS13C to damaged lysosomes precedes mobilization of OSBP and does not require a functional PITT pathway is consistent with recent findings by Wang *et al.*^23^ and prompted us to address a potential role for VPS13C-mediated lipid transport in lysosomal repair.

### VPS13C is essential for tethering damaged lysosomes to the ER

Activation of the PITT pathway initiates lysosomal recruitment of OSBP and multiple OSBP-related protein (ORP) family members to orchestrate extensive new membrane contacts between damaged lysosomes and the ER^21,27^. Consistent with its proposed bridge-like lipid transport function at ER-lysosome contacts, VPS13C bound to damaged lysosomes simultaneously interacts with the transmembrane ER proteins VAPA and VAPB via a FFAT motif located near its *N*-terminus^23,28^. Since VPS13C recruitment to damaged lysosomes occurs independently of the PITT pathway (**Fig. 2d, e**), we asked whether ER-wrapping of damaged lysosomes is dependent on VPS13C. To address this, U2OS cells co-expressing GFP fused to the *N*-terminal ER membrane anchor of cytochrome P450 and the cytosolic SM reporter EqtSM-Halo were treated with LLOMe, fixed and immunostained for LAMP1. After LLOMe treatment, an extensive wrapping of LAMP1-positive lysosomes with GFP-positive ER membrane was observed, which was accompanied by lysosomal translocation of EqtSM (**Fig. 3a**). Removal of VPS13C or PI4K2A in each case abolished LLOMe-induced ER-lysosome wrapping and resulted in an enhanced EqtSM labeling of damaged lysosomes (**Fig. 3a; Suppl. 2b**). These trends were even more apparent in cells pre-treated with apilimod, a PIKfyve phosphoinositide kinase inhibitor that causes a massive swelling of endolysosomes^29^. In a complementary approach, wildtype, VPS13C-KO and PI4K2A-KO cells were treated with LLOMe in the absence or presence of apilimod and then immunostained for VAPA and LAMP1. Under resting conditions, none of the cell lines showed any appreciable mobilization of VAPA to the LAMP1-positive lysosomes (**Fig. 3b**). However, LLOMe treatment in wildtype cells caused a robust mobilization of VAPA to both control and swollen lysosomes, as evidenced by an extensive co-localization with LAMP1. In contrast, in both independent VPS13C-KO cell lines and PI4K2-KO cells, LLOMe-induced recruitment of VAPA to lysosomes was abolished (**Fig. 3b, c**). Together, these results indicate that both VPS13C and the PITT pathway are essential for expanding ER-lysosome membrane contacts in response to lysosomal damage.

**Figure 3.**
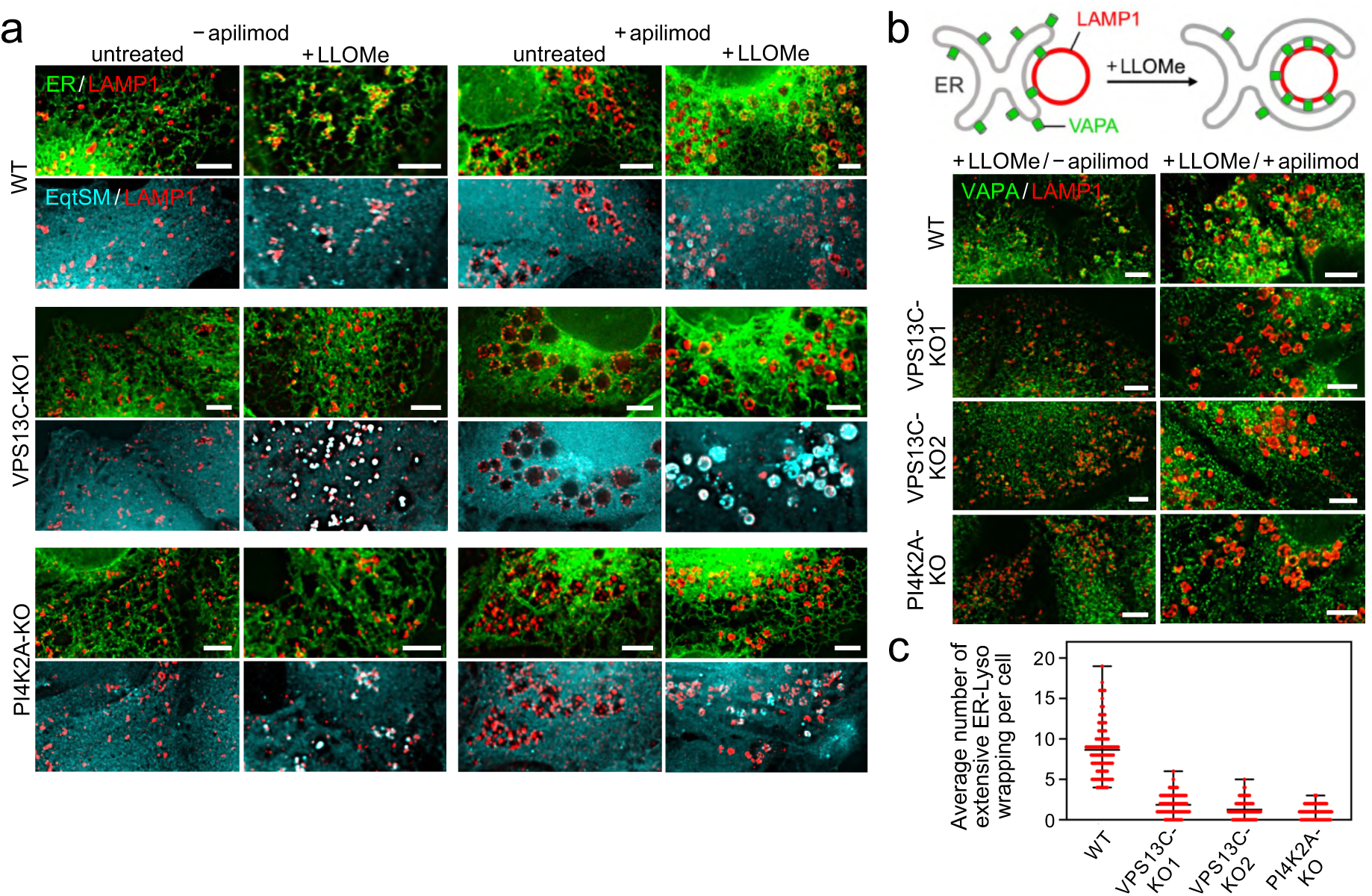
VPS13C is essential for tethering damaged lysosomes to the ER. (**a**) U2OS wildtype (WT), VPS13C-KO1 and PI4K2A-KO cells co-expressing EqtSM-Halo (*cyan*) and GFP fused to the *N*-terminal ER membrane anchor of cytochrome P450 (*green*) were treated with apilimod (200 nM, 2 h) in combination with LLOMe (1 mM, 10 min), immunostained for LAMP1 (*red*) and imaged by DeltaVision microscopy. Scale bar, 5 µm. (**b**) U2OS wildtype (WT), VPS13C-KO1, VPS13C-KO2 and PI4K2A-KO cells were treated with apilimod (200 nM, 2 h) in combination with LLOMe (1 mM, 10 min), immunostained for LAMP1 (*red*) and VAPA (green), and imaged by DeltaVision microscopy. Scale bar, 5 µm. (**c**) Average number of LAMP1-positive lysosomes extensively wrapped with VAPA-positive ER per cell in cells treated with apilimod and LLOMe as in (b). Data are means ± SD (*n* = 105 cells per cell line) from 3 independent experiments.

### VPS13C readily binds lysosomes under mechanical or osmotic stress

Tracing individual lysosomes over time to assess dynamic processes occurring on their surface is challenging owing to their small size, heterogenous morphology, and motility^30^. Recent work showed that non-phagocytic cells take up large (3 μm) polystyrene microbeads that accumulate in acidified compartments^31^. To test whether this approach could be used to enlarge lysosomes and trace them individually over time, we fed U2OS cells with polystyrene microbeads functionalized with the pH-sensitive fluorophore pHrodo SE (**Fig. 4a**). While non-internalized microbeads did not exhibit any pHrodo signal, internalized microbeads readily became fluorescent and localized within LAMP1-positive membranes, indicating delivery to lysosomal compartments (**Fig. 4b; Suppl. Fig. 3**). Within minutes of LLOMe treatment, microbead-containing lysosomes lost their pHrodo fluorescence and simultaneously accumulated the cytosolic SM reporter EqtSM-Halo on their surface (**Fig. 4b**), analogous to the neutralization of acidic pH and SM scrambling observed in damaged native lysosomes^18^.

**Figure 4.**
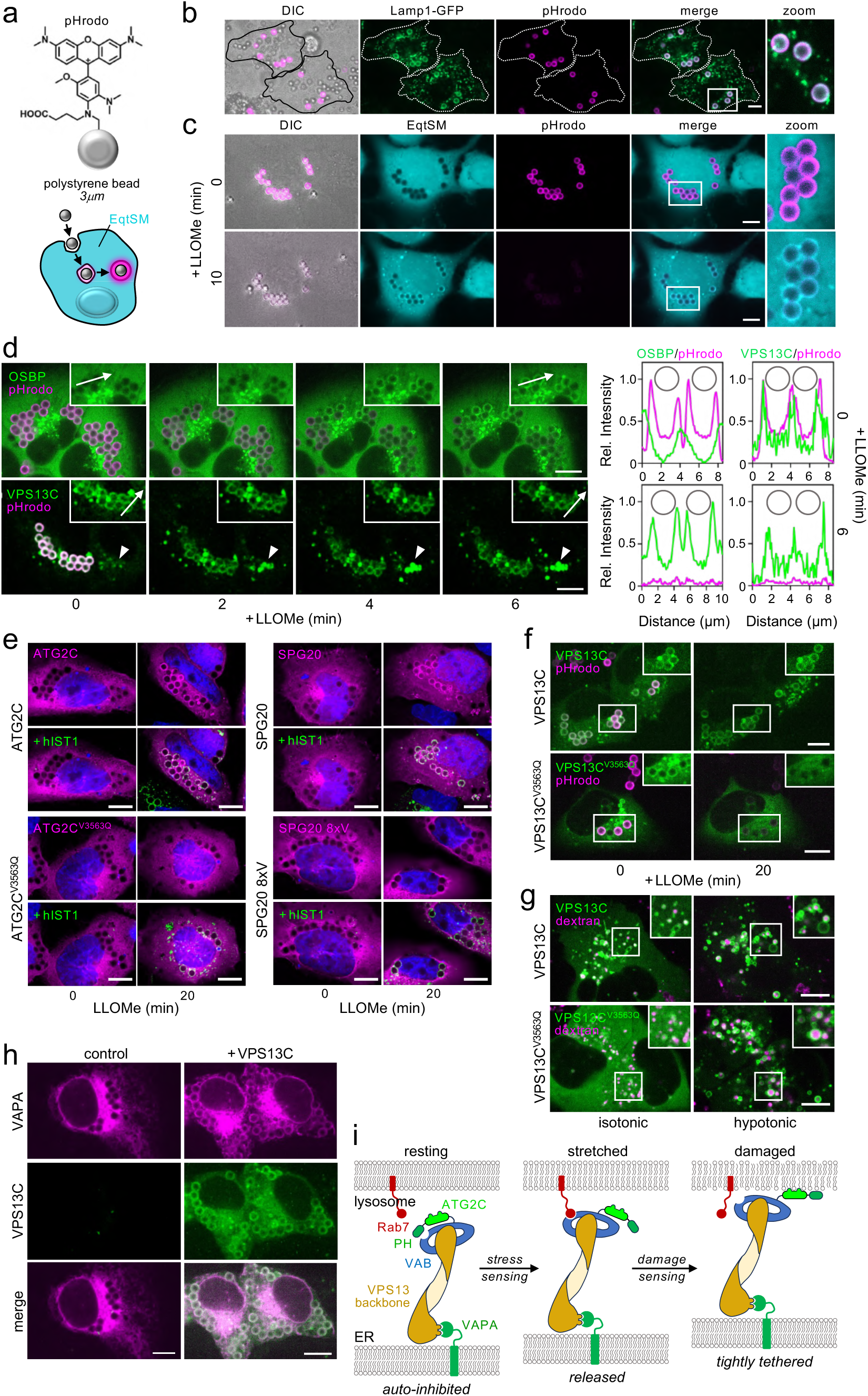
VPS13C readily binds microbead-containing lysosomes. (**a**) Schematic of the cellular uptake of 3 µm polystyrene microbeads functionalized with the pH-sensitive dye pHrodo SE. (**b**) Differential interference contrast (DIC) and fluorescence images of HeLa cells expressing GFP-tagged LAMP1 (*green*) that were fed pHrodo-microbeads (*magenta*) for 24 h. Cells were imaged by spinning disk microscopy. Scale bar, 10 µm. (**c**) Time-lapse DIC and fluorescence images of HeLa cells expressing Halo-tagged EqtSM (*cyan*), fed pHrodo-microbeads (*magenta*) and treated with 1 mM LLOMe for the indicated time. Cells were imaged by spinning disk microscopy. Scale bar, 10 µm. (**d**) Time-lapse fluorescence images of U2OS cells expressing OSBP-GFP or VPS13C-mClover (*green*) that were fed pHrodo-microbeads (*magenta*) and treated with 1 mM LLOMe for the indicated time. Line scans show overlap of OSBP or VPS13C and pHrodo signals along the path of arrow, with the position of microbeads marked by grey circles. Scale bar, 10 µm. (**e**) Fluorescence images of U2OS cells containing 3 µm polystyrene microbeads and expressing ATG2C-Halo, ATG2C^V3563Q^-Halo, SPG20-GFP or SPG20 8xV-GFP (*magenta*) were treated with 1 mM LLOMe for the indicated time, and then immunostained for hIST1 (*green*) and counter stained with DAPI (*blue*). Cells were imaged by DeltaVision microscopy. Scale bar, 10 µm. (**f**) Time-lapse fluorescence images of U2OS cells expressing mClover-tagged VPS13C or VPS13C^V3563Q^ (*green*), fed pHrodo-microbeads (*magenta*) and treated with 1 mM LLOMe for the indicated time. Cells were imaged by spinning disk microscopy. Scale bar, 10 µm. (**g**) Time-lapse fluorescence images of U2OS cells expressing VPS13C-GFP or VPS13C^V3563Q^-GFP (*green*) and fed CF633-labelled 70kDa dextran (*magenta*) were incubated in isotonic medium (100% Opti-MEM) and then incubated for 5 min in hypotonic medium (1% Opti-MEM). Scale bar, 10 µm. (**h**) Live-cell fluorescence images of U2OS VPS13C-KO1 cells that were fed 3 µm polystyrene microbeads and then co-transfected with VAPA-mCherry (*magenta*) and VPS13C-mClover (*green*) as indicated. Scale bar, 10 µm. (**i**) Schematic of how conformational flexibility of its ATG2C domain-containing *C*-terminus may enable VPS13C to distinguish stressed from injured lysosomes. See text for further details. Illustration adapted from^23^.

We next asked whether microbead-containing lysosomes mobilize lipid transport machinery in response to damage. GFP-tagged OSBP, a core component of the PITT pathway^21,27^, was recruited to the surface of microbead-containing lysosomes within minutes after LLOMe addition, concomitant with a loss of pHrodo-fluorescence (**Fig. 4d**). LLOMe-induced mobilization of OSBP was strictly dependent on PI4K2A (**Fig. 5b**), indicating that microbead-containing lysosomes retain the ability to activate the PITT pathway in response to damage. Strikingly, VPS13C-mClover was already massively mobilized to the surface of microbead-containing lysosomes under resting conditions (**Fig. 4d**). Subsequent LLOMe-treatment enhanced the recruitment of VPS13C-mClover to native, but not microbead-containing lysosomes. This suggests that polystyrene microbeads perturb the surrounding lysosomal membrane to a degree sufficient to trigger large-scale VPS13C binding but inadequate for inducing proton leakage or PITT pathway activation.

**Figure 5.**
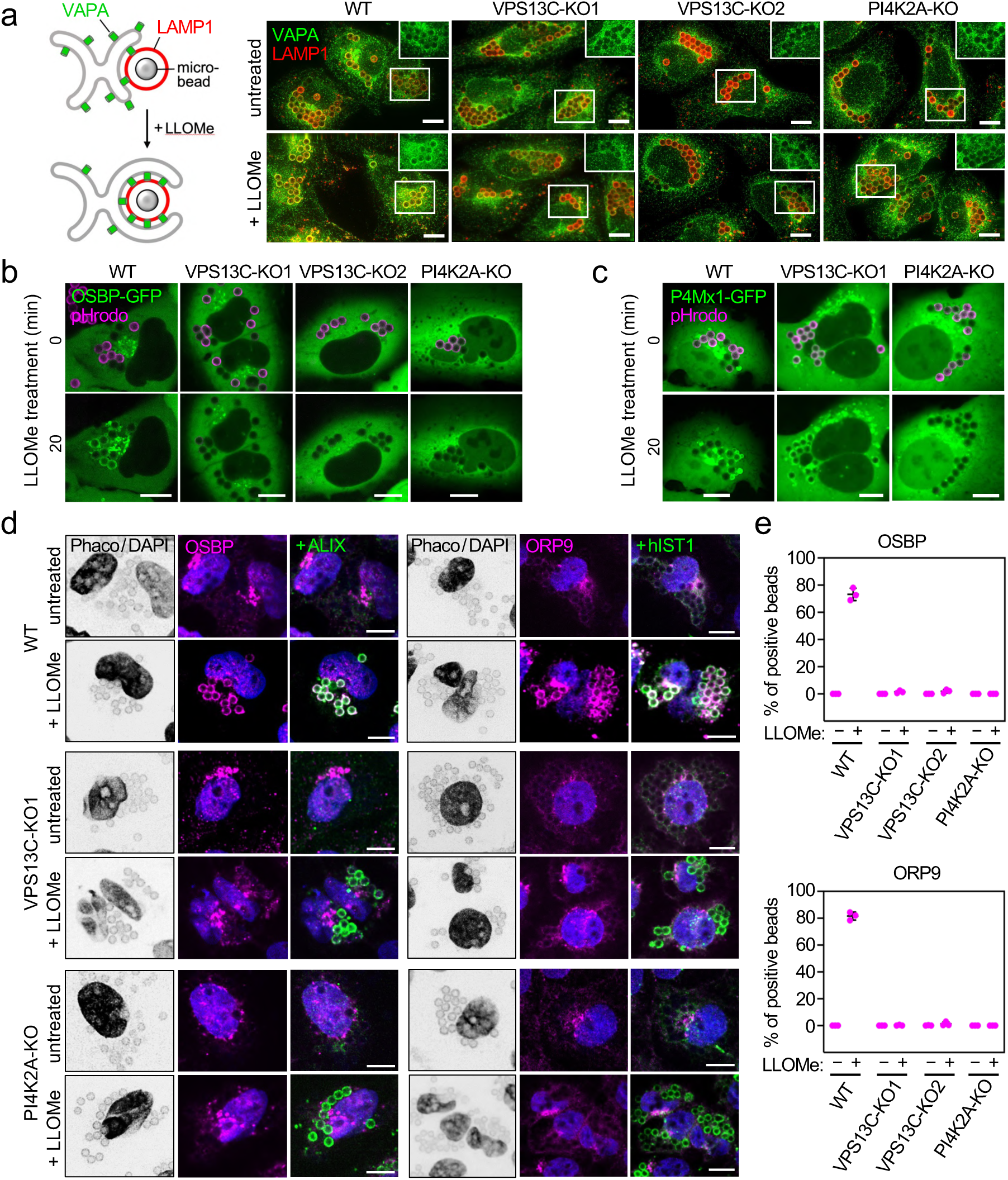
VPS13C is essential for recruiting OSBP and ORP9 to damaged lysosomes. (**a**) U2OS wildtype (WT), VPS13C-KO1, VPS13C-KO2 or PI4K2A-KO cells were fed 3 μm polystyrene microbeads, treated with LLOMe (1 mM, 10 min) as indicated, immunostained for LAMP1 (*red*) and VAPA (*green*), and imaged by DeltaVision microscopy. Scale bar, 10 µm. (**b**) Time-lapse images of U2OS wildtype (WT), VPS13C-KO1, VPS13C-KO2 or PI4K2A-KO cells containing 3 µm pHrodo-labeled microbeads (*magenta*), expressing GFP-tagged OSBP (*green*) and treated with 1 mM LLOMe for the indicated time. Cells were imaged by spinning disk microscopy. Scale bar, 10 µm. (**c**) Time-lapse images of U2OS wildtype (WT), VPS13C-KO1 or PI4K2A-KO cells containing 3 µm pHrodo-labeled microbeads (*magenta*), expressing GFP-tagged PI4P sensor P4MX1 (*green*) and treated with 1 mM LLOMe for the indicated time. Cells were imaged by spinning disk microscopy. Scale bar, 10 µm. (**d**) Phase contrast (Phaco) and fluorescence images of polystyrene microbead-containing untreated or LLOMe-treated (1 mM, 10 min) U2OS wildtype (WT), VPS13C-KO1 or PI4K2A-KO cells immunostained for OSBP (*magenta*) and ALIX (*green*) or ORP9 (*magenta*) and hIST1 (*green*) and counterstained with DAPI (*blue*). Cells were imaged by spinning disk microscopy. Scale bar, 10 µm. (**e**) Percentage of microbead-containing lysosomes with positive immunostaining for OSBP or ORP9 in cells treated as in (d). Data are means ± SD of three independent experiments.

The *C*-terminal region of VPS13C harbors an ATG2C domain and a PH domain, which project out of the rod-like core of the protein^32^. The ATG2C domain comprises a bundle of four amphipathic helices that confers affinity of the protein for lipid droplets, presumably owing to packing defects in the phospholipid monolayer that surrounds them^28^. We considered that polystyrene microbeads inside lysosomes may trigger binding of VPS13C via its ATG2C domain through recognition of lipid packing defects caused by membrane stretching. To test this, we expressed a Halo-tagged version of ATG2C_VPS13C_ in bead-containing U2OS cells. As control, we also expressed GFP-tagged SPG20/SPART, a protein mobilized to the surface of damaged lysosomes via sensory amphipathic helices that recognize damage-associated lipid packing defects^4,33^. SPG20 did not bind microbead-containing lysosomes unless cells were treated with LLOMe (**Fig. 4e**). LLOMe-induced recruitment of SPG20 was abolished when 8 bulky hydrophobic residues in its amphipathic helices were changed to the small hydrophobic valine (SPG20 8xV). Unlike full-length VPS13C, ATG2C_VPS13C_ binding to microbead-containing lysosomes occurred exclusively upon treatment with LLOMe. Mutation of a single residue (V3563Q) on the hydrophobic face of the amphipathic helix bundle in ATG2C_VPS13C_ was sufficient to abolish its LLOMe-induced recruitment to microbead-containing lysosomes (**Fig. 4e**).

Strikingly, a full-length VPS13C variant with the same mutation was efficiently recruited to microbead-containing lysosomes under resting conditions but redistributed to the cytosol upon LLOMe treatment (**Fig. 4f**). This indicates that an intact ATG2C_VPS13C_ domain is essential to mediate VPS13C binding to damaged lysosomes but dispensable for VPS13C binding to lysosomes under mechanical stress/tension. Consistent with this idea, we observed that hypotonic swelling of lysosomes induced a massive mobilization of both VPS13C and VPS13C^V3563Q^ on their surface (**Fig. 4g**).

Wang *et al*.^23^ previously reported that VPS13C can bind the lysosomal GTPase Rab7 via its VAB domain but that intramolecular interactions involving its *C*-terminal ATG2C_VPS13C_ domain hinder the VAB domain from assessing Rab7 under resting conditions. The authors proposed that membrane packing defects induced by lysosomal damage promote membrane binding of the ATG2C_VPS13C_ domain, causing a conformational change in the *C*-terminal region of VPS13C that releases the VAB domain and enables its interaction with Rab7, thereby enhancing binding of VPS13C to lysosomes. However, our data suggest that release of the VAB domain and enhanced Rab7 binding of VPS13C may already be triggered by lysosomes under mechanical or hypoosmotic tension prior to the manifestation of damage-induced lipid packing defects (**Fig. 4i**). While the underlying mechanism remains to be established, it appears that stretching the lysosome-limiting membrane by microbeads or osmotic pressure is sufficient to initiate large-scale mobilization of VPS13C on the lysosomal surface without the need for a functional ATG2C domain. However, in response to lysosome damage, binding of VPS13C to lysosomes becomes critically dependent on its ATG2C domain. These results indicate that recognition of stressed and damaged lysosomes by VPS13C is mediated by distinct mechanisms and relies on different protein conformations (**Fig. 4i**). Our findings reinforce the notion that VPS13C acts upstream of the PITT pathway in response to lysosomal damage.

### Loss of VPS13C disrupts OSBP and ORP9 recruitment to damaged lysosomes

We observed that expression of VPS13C-mClover in U2OS VPS13C-KO cells caused a robust mobilization of co-expressed VAPA-mCherry to microbead-containing lysosomes (**Fig. 4h**). This suggested that mechanically stressed lysosomes may already induce VPS13C-mediated ER wrapping even in the absence of membrane damage. To further dissect the role of VPS13C and the PITT pathway in the formation of ER contacts with mechanically stressed versus damaged lysosomes, wildtype, VPS13C-KO and PI4K2A-KO cells were fed microbeads and then immunostained for endogenous VAPA and LAMP1 before and after LLOMe treatment. Under resting conditions, none of the cell lines showed any appreciable mobilization of VAPA to the microbead-containing lysosomes (**Fig. 5a; Suppl. Fig. 4**). However, LLOMe treatment in wildtype cells caused a robust recruitment of VAPA to microbead-containing lysosomes, as evidenced by extensive co-localization with LAMP1. In contrast, both VPS13C-KO and PI4K2A-KO cells failed to accumulate VAPA on the surface of microbead-containing lysosomes in response to LLOMe.

The foregoing implied that VPS13C on its own is not sufficient to mediate large-scale VAPA recruitment to damaged lysosomes and that efficient ER wrapping relies on the mobilization of OSBP/ORP family lipid transport proteins via the PITT pathway^21^. As VPS13C recruitment to damaged lysosomes precedes mobilization of OSBP (**Fig. 2e**), we asked whether binding of OSBP to damaged lysosomes requires VPS13C. Using two independent VPS13C-KO cell lines, we observed that VPS13C removal completely abolished LLOMe-induced recruitment of GFP-OSBP to microbead-containing lysosomes (**Fig. 5b**). As OSBP binds damaged lysosomes via PI4P produced by PI4K2A on their surface^21,27^, we asked whether VPS13C is essential for the damage-induced accumulation of lysosomal PI4P. In VPS13C-KO cells treated with LLOMe, we observed a robust recruitment of the GFP-tagged PI4P biosensor P4XM1 to microbead-containing lysosomes (**Fig. 5c**). In contrast, P4Mx1-GFP remained entirely cytosolic in LLOMe-treated PI4K2A-KO cells. We then analyzed the impact of VPS13C and PI4K2A removal on the LLOMe-induced mobilization of endogenous OSBP and ORP family member ORP9 to microbead-containing lysosomes by immunofluorescence microscopy. This revealed that binding of OSBP and ORP9 to damaged lysosomes in each case required both VPS13C and PI4K2A (**Fig. 5d, e; Suppl. Fig. 5**). In contrast, VPS13C removal had no impact on LLOMe-induced mobilization of PI4K2A (**Suppl. Fig. 6**), in line with our finding that VPS13C is dispensable for the damage-induced accumulation of PI4P on lysosomes. Together, these data indicate that VPS13C is required for the mobilization of OSBP, ORP9 and potentially other ER-lysosome tethers to damaged lysosomes to enable their efficient ER wrapping.

### Lysosomal damage induces directional ER-to-lysosome lipid transport

The recruitment of VPS13C to membrane contact sites between the ER and damaged lysosomes has been proposed to create a path for bulk lipid transport from the ER to lysosomes to enable efficient lysosomal repair^23^. To experimentally address this scenario, we adopted a chemical strategy previously used to image ER-derived lipid flows in living cells in real time^34^. This method relies on the metabolic incorporation of azido-choline (N_3_-Cho) into the cellular phosphatidylcholine (PC) pool, followed by spatially limited strain-promoted alkyne–azide cycloaddition (SPAAC) of an ER-localizable fluorophore bearing a strained alkyne, BDP-FL-DBCO (ER-DBCO; **Fig. 6a**). Box-like LTPs like OSBP and ORP9 typically select one lipid at a time for transport based on headgroup recognition. Consequently, they would likely fail to accommodate lipids with DBCO-modified head groups. In contrast, bridge-like LTPs such as VPS13C are more permissive, as they provide a long hydrophobic groove that can accommodate the acyl chains of multiple lipids at a time to enable their bulk transport with minimal head group recognition^35,36^. Moreover, an energy minimization model revealed that the headgroup of PC containing a modified cyclooctane ring system derived from DBCO retains substantial conformational flexibility after the SPAAC reaction (**Fig. 6b**), in line with the plasticity reported for related fused ring systems^37^. This inherent flexibility of the cyclooctane ring may facilitate passage of DBCO-modified PC molecules through the narrow tunnel of bridge-like LTPs^38^.

**Figure 6.**
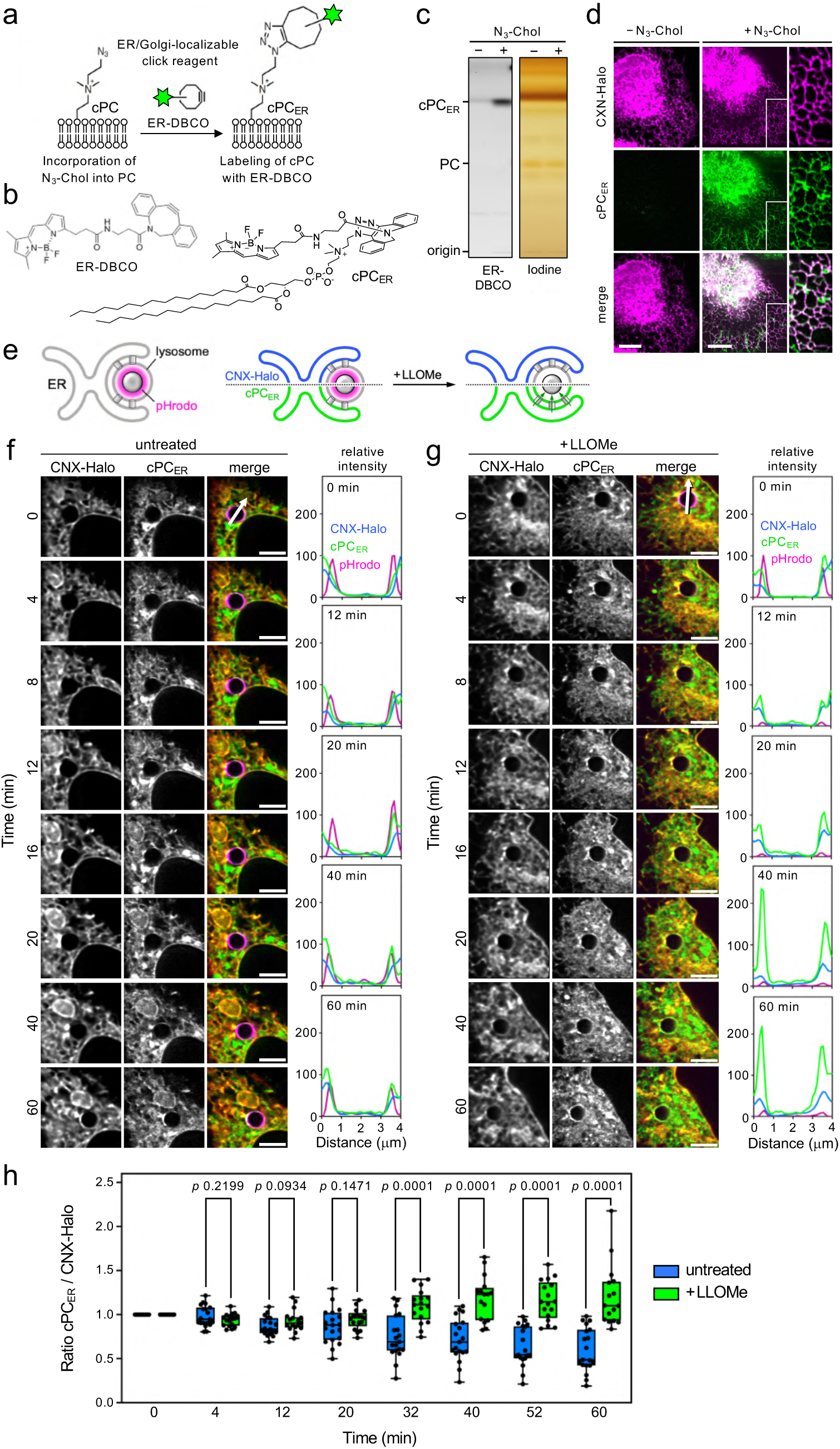
Lysosomal damage promotes ER-to-lysosome lipid transport. (**a**) Schematic illustrating metabolic incorporation of N_3_-choline (N_3_-Chol) into the cellular phosphatidylcholine (PC) pool followed by labeling of the N_3_-choline head with the ER/Golgi-targeted fluorescent click reagent BDP-FL-DBCO (ER-DBCO). (**b**) Chemical structure of click reagent ER-DBCO and energy minimization model of an ER-DBCO click-reacted PC molecule (cPC_ER_). An envelope conformation of the cyclooctane ring in the DBCO-moiety of cPC_ER_ represents the lowest energy and thereby most stable conformation. Energy minimization model created with the Chem3D tool of ChemDraw (version 23.0). (**c**) HeLa cells were cultured in the presence or absence of N_3_-Chol (24 h, 500 μM), washed, stained with ER-DBCO (15 min, 500 μM), washed, incubated for 30 min and subjected to lipid extraction and TLC analysis. Total lipids were stained using iodine vapor. (**d**) U2OS cells expressing Halo-tagged calnexin (CNX-Halo, *magenta*) were cultured in the presence or absence of N_3_-Chol (24 h, 500 μM), washed, stained with ER-DBCO (15 min, 500 μM, *green*), washed, incubated for 30 min and imaged by lattice light-sheet microscopy (LLSM). Scale bar, 10 µm. (**e**) Schematic of ratiometric approach used to detect bulk lipid transport from the ER to damaged lysosomes containing pHrodo microbeads. See text for details. (**f**) U2OS cells expressing CNX-Halo (*red*) were fed pHrodo-microbeads (*magenta*), incubated with N_3_-Chol, stained with ER-DBCO as in (d) (cPC_ER_, *green*) and then subjected to time-lapse imaging by LLSM. Line scans show the intensity profiles of CNX-Halo, pHrodo and cPC_ER_ signals along the path of the arrows. Profiles are plotted as relative intensities for each channel normalized to the 0 min time point. Scale bar, 5 µm. (**g**) Time-lapse images and line scans of U2OS cells as in (g) but treated with 1 mM LLOMe for the indicated time points. Scale bar, 5 µm. (**h**) Ratiometric analysis of cPC_ER_ and CNX-Halo fluorescence levels on the surface of pHrodo-microbead containing lysosomes in untreated and LLOMe-treated cells. For each individual microbead, the relative change in cPC_ER_ and CNX-Halo signal was traced over time. To assess the relative enrichment of cPC_ER_ over that of CNX-Halo, the change in cPC_ER_ fluorescence relative to the 0 min time point was divided by the corresponding change in CNX-Halo fluorescence to obtain a ratio that reflects differential signal development over time. Data shown are cPC_ER_/CNX-Halo ratios after normalization against the 0 min time point value, which was set at 1.0. *P*-values were calculated using an unpaired two-tailed Student’s *t*-test with Welch’s correction. Control, *n* = 19 microbeads from 10 different cells; + LLOMe: *n* = 17 microbeads from 8 different cells, each from at least 3 independent measurements.

TLC analysis showed that ER-DBCO selectively click-reacted with PC in cells metabolically labeled with N_3_-Cho (**Fig. 6c**). This reaction did not occur when N_3_-Cho was omitted. Live cell imaging of N_3_-Cho-labeled U2OS cells revealed extensive co-localization of ER-DBCO fluorescence with the Halo-tagged ER membrane protein calnexin (CXN-Halo; **Fig. 6d**), indicating that ER-DBCO primarily reacted with clickable PC in the ER (hereafter referred to cPC_ER_). No ER-DBCO signal was observed when N_3_-Cho was omitted. Having established selective labeling of an ER-resident PC pool under live-cell conditions, we next developed a method to quantitatively assess ER-derived PC transport to damaged lysosomes in real time. As ER-lysosome contact sites bridged by VPS13C have an estimated gap width of ∼33.5 nm^32^, using light microscopy to distinguish the ER from the lysosomal membrane at these sites is not possible. An additional complexity is that lysosomal damage causes extensive ER-wrapping of the injured organelle. To circumvent these challenges, we pursued a ratiometric approach in which we simultaneously traced the fluorescence levels of cPC_ER_ and CNX-Halo on the surface of lysosomes containing pHrodo-labelled microbeads by lattice light-sheet microscopy (LLSM). The rationale behind this approach was that, while damage-induced ER wrapping would simultaneously amplify the cPC_ER_ and CNX-Halo signals around the microbead-containing lysosomes, and therefore not alter their ratio, bulk transport of cPC_ER_ to the lysosome would gradually enhance its signal relative to that of CNX-Halo (**Fig. 6e**).

Under resting conditions, the pHrodo signal of the microbead-containing lysosomes was retained while no increase in either cPC_ER_ or CNX-Halo signal on their surface was observed during 60 min of imaging (**Fig. 6f**). In cells treated with LLOMe, the pHrodo signal readily disappeared, as expected. This was accompanied by a moderate increase in both cPC_ER_ and CNX-Halo signals within the first 20 min of imaging (**Fig. 6g**). The latter was evident by a more pronounced ring-like staining around the microbead-containing lysosomes, indicative of enhanced ER wrapping. Also, sharper peaks in the corresponding line scan plots, representing ER membrane around the microbeads, supported this observation. However, after prolonged incubation (>20 min), cPC_ER_ fluorescence surrounding the microbeads increased substantially while the CNX-Halo signal remained largely unchanged relative to earlier timepoints. To quantitatively assess the enrichment in cPC_ER_ fluorescence over that of CNX-Halo, the change in cPC_ER_ fluorescence relative to the 0 min time point was divided by the corresponding change in CNX-Halo fluorescence to obtain a ratio that reflects differential signal development over time. To enable a comparative analysis of changes in fluorescence levels among individual microbead-containing lysosomes across different experiments, the cPC_ER_/CNX-Halo ratio measured at the 0 min time point was set at 1.0. In cells under resting conditions, the cPC_ER_/CNX-Halo ratio gradually decreased from 1.0 to 0.6 over a time period of 60 min (**Fig. 6h**). This drop in ratio value could be due to photobleaching of cPC_ER_ and/or its transport out of the ER to other organelles, as previously described^34^. In LLOMe-treated cells, on the other hand, the cPC_ER_/CNX-Halo ratio remained close to 1.0 during the first 20 min of imaging and then gradually exceeded 1.0 at later time points, reaching a value twice as high as that measured in resting cells after 60 min of imaging (**Fig. 6h**). Collectively, these results indicate that the LLOMe-induced expansion of ER-lysosome contact sites is accompanied by net transport of cPC_ER_ to the lysosomal membrane.

### Delivery of ER lipids to damaged lysosomes requires VPS13C

Having shown that lysosomal damage induces directional ER-to-lysosome lipid transport, we next asked whether this process relies on VPS13C. To address this, we used cells stably transduced with EqtSM-Halo as additional marker of lysosomal damage. In wildtype cells under resting conditions, microbead-containing lysosomes retained pHrodo signal over 60 min of imaging without displaying any appreciable increase in cPC_ER_ or EqtSM fluorescence on their surface (**Fig. 7a; Suppl. Fig. 7a**). In contrast, addition of LLOMe readily abolished pHrodo fluorescence concomitant with an instant rise in both EqtSM and cPC_ER_ signals around the microbead-containing lysosomes at early timepoints (4-20 min). While the EqtSM signal reached a plateau at prolonged imaging times, the rise in cPC_ER_ signal persisted over the entire 60 min imaging period, consistent with our previous findings (**Fig. 7a; Suppl. Fig. 7a**). In two independent VPS13C-KO cell lines under resting conditions, no change in either EqtSM, cPC_ER_ or pHrodo signal around the microbead-containing lysosomes was seen (**Fig. 7b; Suppl. Fig. 7b**). Following LLOMe treatment, EqtSM displayed an instant and robust labeling of microbead-containing lysosomes, which was notably more pronounced than that seen in wildtype cells. At the same time, cPC_ER_ accumulation was completely abolished in both VPS13C-KO cell lines (**Fig. 7b; Suppl. Fig. 7b, c**). Quantification of fluorescence signals of the different channels on 3D-surfaces defined by pHrodo-microbeads imaged in cells across multiple independent experiments confirmed these observations (**Fig. 7c**). In contrast, removal of ATG13, a protein essential for autophagy initiation^39–41^, had no impact on cPC_ER_ accumulation on damaged lysosomes (**Suppl. Fig. 8a, b**), indicating that this process occurs independently of autophagosome formation. Collectively, these results demonstrate that VPS13C is essential for expanding ER membrane contacts with damaged lysosomes as well as for establishing pipelines that mediate bulk transport of lipids to the lysosome-limiting membrane. While our data do not provide conclusive evidence that VPS13C itself is the transporter responsible for this damage-induced lipid flow, they show that the protein plays an indispensable role.

**Figure 7.**
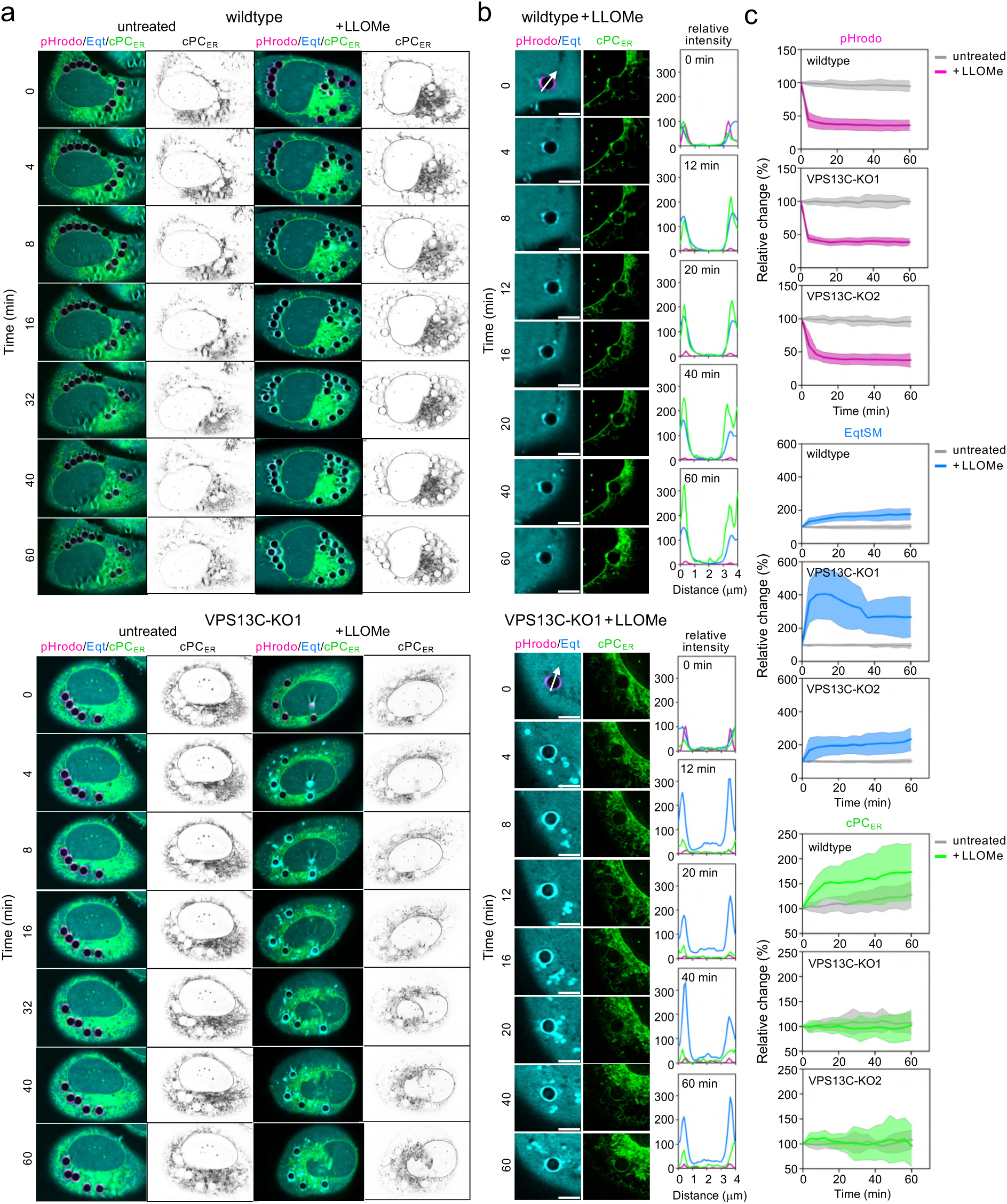
Delivery of ER lipids to damaged lysosomes requires VPS13C. (**a**) U2OS wildtype and VPS13C-KO1 cells expressing EqtSM-Halo (*cyan*) were fed pHrodo-microbeads (*magenta*), incubated with N_3_-Chol, stained with ER-DBCO (cPC_ER_, *green*) and then subjected to time-lapse imaging in the absence (untreated) or presence of 1 mM LLOMe. Cells were imaged by LLSM. (**b**) Line scans showing the intensity profiles of EqtSM-Halo (*cyan*), pHrodo (*magenta*) and cPC_ER_ (*green*) signals along the path of the arrows in the zoom-in of cells treated as in (a). Profiles are plotted as relative intensities for each channel normalized to the 0 min time point. Scale bar, 5 µm. (**c**) Time course plotting the relative changes in pHrodo, EqtSM-Halo, and cPC_ER_ signals on the surface of pHrodo-microbead containing lysosomes in U2OS wildtype, VPS13C-KO1 and VPS13C-KO2 cells treated as in (a). A 3D-surface was generated around the pHrodo-positive microbeads using Imaris software, and the mean fluorescence intensities of the respective channels on this surface were quantified. For each cell, signals from all microbead-associated surfaces were averaged. Values were normalized to the signal at timepoint 0 min and plotted as mean ± SD over time. Wildtype (*n* = 8): control, 19 cells; +LLOMe, 20 cells. VPS13C-KO1 (*n* = 4): control, 10 cells; +LLOMe, 11 cells. VPS13C-KO2 (*n* = 3): control, 9 cells; +LLOMe, 13 cells.

### VPS13C removal disrupts lysosomal repair

Cells lacking VPS13C display an enhanced lysosomal fragility as revealed by a chronic activation of the cGAS-STING pathway of innate immunity^42^ along with an enhanced accumulation of Gal3^23^ and robust cytosolic exposure of SM in response to lysosome-damaging agents (**Fig. 7a-c**). To address whether VPS13C participates in the repair of acutely damaged lysosomes, we took advantage of the lysosomotropic compound glycyl-L-phenylalanine 2-naphtylamide (GPN). In the lysosomal lumen, GPN is processed into metabolites that transiently disrupt lysosomal integrity by promoting osmotic rupture of the lysosome-limiting membrane^43^. As lysosomal pH critically relies on the structural integrity of lysosome-limiting membrane, the process of lysosomal damage and repair can be monitored in real time using lysosomal fluorescent pH sensors such as Lyso-Tracker or Lyso-pHluorin (**Fig. 8a**). While Lyso-Tracker is fluorescent at acidic pH^44^, the fluorescence of Lyso-pHluorin is quenched at acidic pH and activated upon pH neutralization^45^. Consequently, wildtype cells briefly exposed to GPN lose Lyso-Tracker fluorescence and concomitantly accumulate Lyso-pHluorin-positive puncta (**Fig. 8b, c**). When GPN is washed away, Lyso-Tracker fluorescence readily returned while Lyso-pHluorin puncta disappeared quickly, indicative of efficient lysosomal repair. However, in two independent VPS13C-KO cell lines, the recovery of Lyso-Tracker fluorescence and quenching of Lyso-pHluorin fluorescence after GPN removal were strongly impaired (**Fig. 8b, c; Suppl. Fig. 9**). In a complementary set of experiments, we quantitatively assessed the recovery of lysosomes from a brief LLOMe treatment using Lyso-pHluorin as lysosomal damage marker. Lyso-pHluorin-positive puncta were not detected in either wildtype or VPS13C-KO cells under resting conditions. Upon 10 min exposure to LLOMe, most cells displayed numerous Lyso-pHluorin-positive puncta, suggesting equal lysosomal damage (**Fig. 8d**). While Lyso-pHluorin-positive puncta in wildtype cells were readily turned-over, puncta clearance was significantly slower in the two independent VPS13C-KO cell lines (**Fig. 8d, e**). Together, these data demonstrate that VPS13C is essential for efficient repair of acutely damaged lysosomes.

**Figure 8.**
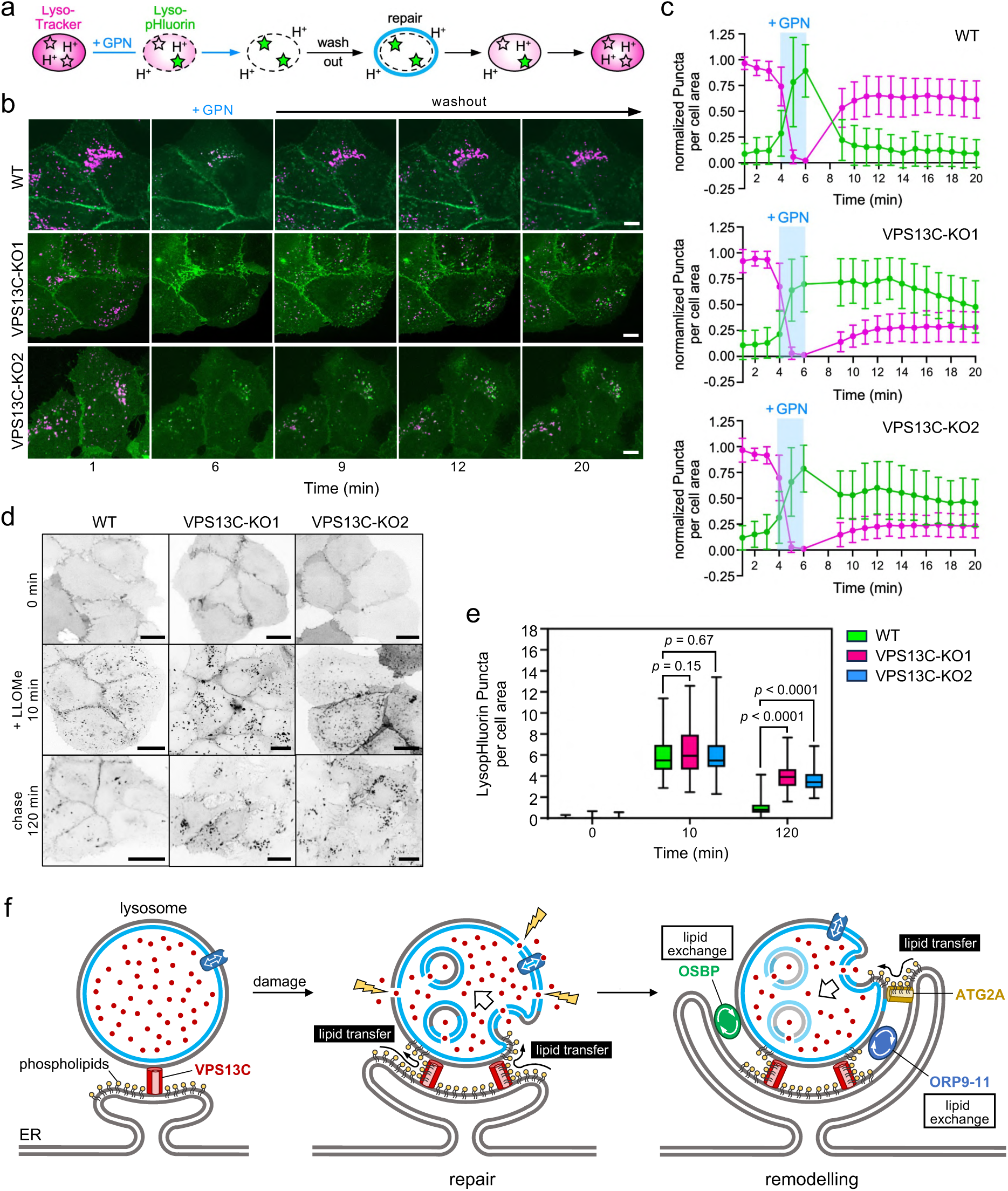
Repair of damaged lysosomes is critically dependent on VPS13C. (**a**) Schematic of lysosomal repair assay based on the use of endolysosomal pH sensors Lyso-Tracker and Lyso-pHluorin. (**b**) Time-lapse images of LysoTracker-labelled (*magenta*) and Lyso-pHluorin-expressing (*green*) U2OS wildtype (WT), VPS13C-KO1 and VPS13C-KO2 cells pulse-treated with GPN (200 μM, 2 min). Cells were imaged by spinning disk microscopy. Scale bar, 10 µm. (**c**) Time-course plotting Lyso-Tracker- and Lyso-pHluorin-positive puncta in cells treated as in (b). Data are means ± SD (*n* = 84 cells for WT, 105 cells for VPS13C-KO1, 86 cells for VPS13C-KO2) from at least 2 independent experiments. (**d**) Time-lapse fluorescence images of Lyso-pHluorin-expressing U2OS wildtype (WT), VPS13C-KO1 and VPS13C-KO2 cells under resting conditions (0 min), after pulse-treatment with LLOMe (1 mM) for 10 min, and after a 120 min chase. Cells were imaged by spinning disk microscopy. Scale bar, 10 µm. (**e**) Time-course plotting Lyso-pHluorin-positive puncta in cells treated as in (d). Data are means ± SD (*n* = 130 cells for WT, 107 cells for VPS13C-KO1 and 93 cells for VPS13C-KO2) from at least 2 independent experiments. (**f**) Schematic illustrating how VPS13C/PARK23 initiates lipid transfer and membrane remodeling to enable efficient lysosomal repair. See text for details.

## DISCUSSION

Our study uncovered an unprecedented role for the bridge-like lipid transport protein VPS13C in sensing lysosomal membrane tension in anticipation of membrane damage to initiate large-scale directional ER-to-lysosome lipid transfer for efficient lysosomal repair. We envision that this damage-induced VPS13C-dependent lipid flow provides lysosomes with bulk lipids from the ER to support formation of intraluminal vesicles, a process crucial for clearing lesions from the lysosome-limiting membrane^15,16,18^. In line with this idea, we observed that cells lacking VPS13C are severely limited in their ability to repair acutely damaged lysosomes. Our findings extend previous studies indicating that loss of VPS13C enhances lysosomal fragility^23,42^ and strengthen the notion that loss-of-function mutations in *VPS13C* increase the risk of Parkinson’s disease by disrupting an effective early lysosomal damage response in cell types that are particularly sensitive to lysosome dysfunction.

Lysosomal damage triggers phosphoinositide signalling via the PITT pathway to mobilize ER-bound OSBP/ORP lipid transport proteins that collectively mediate an expansion of ER-lysosome membrane contact sites in a process that is dependent on the damage-induced production of PI4P on the lysosomal surface^21,27^. OSBP and ORPs subsequently catalyse ER-to-lysosome transfer of cholesterol and PS in exchange for lysosome-derived PI4P. Although this counter lipid transport does not result in net lipid transfer to lysosomes, ORP family-mediated accumulation of PS on the lysosome surface stimulates delivery of bulk lipids for lysosomal repair by the bridge-like lipid transport protein ATG2A^21^. In addition, OSBP-mediated supply of cholesterol to the lysosomal membrane may facilitate net lipid transfer to lysosomes by correcting interleaflet lipid imbalance^46,47^. Convergent with the findings of Wang *et al*.^23^, our data indicate that VPS13C recruitment to damaged lysosomes occurs independently of the PITT pathway and precedes the mobilization of OSBP. Strikingly, loss of VPS13C abolished recruitment of OSBP/ORP lipid transport proteins, VAPA clustering and ER wrapping in response to lysosomal damage. However, lysosomal translocation of PI4K2A and the accumulation of PI4P on the lysosomal surface remained largely unaffected. Collectively, these findings indicate that ER-lysosome contacts formed by VPS13C serve as critical binding platforms for OSBP and ORPs to enable efficient ER wrapping of damaged lysosomes. Moreover, a chemical approach to quantitatively assess directional ER-to-lysosome lipid transport in real time revealed that VPS13C is essential for net delivery of bulk lipids to damaged lysosomes. While our data do not provide ultimate proof that VPS13C is directly responsible for high-volume lipid transport to support lysosomal repair, they unequivocally demonstrate that VPS13C is indispensable for orchestrating a large-scale ER-to-lysosome lipid transfer and acts upstream of OSBP/ORP/ATG2A lipid transport proteins to help preserve lysosome integrity (**Figure 8f**).

The notion that VPS13C is an integral part of an early protective response to lysosomal damage is reinforced by our finding that the protein readily binds to lysosomes under mechanical or hypotonic tension, a condition in which the lysosome-limiting membrane is stretched but not leaky for protons or Ca^2+^. A recent proteome-wide screen for proteins mobilized to osmotically stressed lysosomes yielded both VPS13C and the ER-anchored bridge-like lipid transfer protein LYVAC^47^. Unlike VPS13C, LYVAC proved to be essential for lysosomal vacuolation. Conversely, loss of LYVAC did not impair lysosomal repair^47^, whereas this process was severely perturbed in cells lacking VPS13C (this study). While VPS13C and LYVAC appear to share a common role in lysosomal stress sensing, lipid transport by LYVAC is mainly dedicated to facilitate membrane expansion in order to enhance lysosomal osmoresilience and prevent rupture^47^. In contrast, our data indicate that VPS13C-initiated lipid transfer is primarily linked to the clearance of membrane lesions from damaged lysosomes.

VPS13C recruitment to damaged lysosomes requires release of an auto-inhibited state of the protein, which hinders its access to lysosome-bound Rab7^23^. It was proposed that this conformational switch is triggered by membrane damage-induced packing defects that are recognized by amphipathic helices located in the *C*-terminal ATG2C domain of VPS13C. Interestingly, we observed that mutation of a key residue on the hydrophobic face of a critical amphipathic helix in ATG2C_VPS13C_ abolished binding of VPS13C to damaged lysosomes without affecting its large-scale mobilization to mechanically or osmotically stressed lysosomes. This implies that release of the auto-inhibited state and lysosome binding of VPS13C is part of a lysosomal stress response pathway that does not rely on recognition of lipid packing defects by the ATG2C_VPS13C_ domain. While the underlying mechanism remains to be established, we postulate that binding of VPS13C to stressed lysosomes serves to prime cells for rapid lysosomal repair in anticipation of potential membrane damage. Our data indicate that lipid packing defects induced by lysosomal damage trigger a second conformational change in VPS13C whereby lysosome binding becomes strictly dependent on its ATG2C_VPS13C_ domain (**Fig. 4i**). This second conformational switch may serve to bring the *C*-terminal end of the RGB domain, which forms the central hydrophobic groove of the protein, close to the lysosomal surface, allowing lipids to flow into the lysosome-limiting membrane. In support of this notion, in silico modelling studies revealed that membrane anchoring of the ATG2A domain of VPS13 family members plays a crucial role in conducting lipid transfer to the target membrane^36,48^. The conformational flexibility of its *C*-terminus combined with its ability to distinguish injured from stressed lysosomes makes VPS13C ideally suited to initiate dynamic cycles of directional lipid transfer in synchrony with lysosomal damage and repair, thereby minimizing the risk of lipid back flow.

## METHODS

### Chemical reagents

Chemical reagents were used at the following concentrations, unless indicated otherwise: 1 mM L-leucyl-L-leucine O-methyl ester (LLOMe; Bachem, 4000725), 200 µM glycyl-L-phenylalanine 2-naphtylamide (GPN; Abcam, ab145914), 75 nM LysoTracker^TM^ Red DND-99 (Thermo Fisher Scientific; L7528), 200 nM Apilimod (Sigma-Aldrich, SML2974), 1.7 µM CF®633 dextran 70,000 MW (Biotium, 80141), 100 µg/ml 3 µm Polybead® Amino Microspheres (Polysciences Inc., 17145-5), 100 mM pHrodo^TM^ Red Succinimidylester (ThermoFisher, P36600) and 100 nM BDP-FL-DBCO (Broadpharm, BP-23473). Azido-choline (N_3_-Cho) was synthesized as described in Jao *et al.*^49^ and used at a final concentration of 500 μM.

### Antibodies

Antibodies used were: rabbit polyclonal anti-hIST1 (Proteintech, 19842; IB: 1:1000; IF 1:1000), rabbit polyclonal anti-OSBP (Proteintech, 11096-1-AP; IB: 1:500; IF: 1:200), mouse monoclonal anti-PI4-kinase II Alpha (B5) (Santa Cruz, sc-390026; IB 1:1000; IF 1:400), rabbit polyclonal anti-VPS13C (Proteintech, 28676-1-AP; IB 1:1000), mouse monoclonal anti-α-Actin (Sigma, A1978; IB 1:50,000), mouse monoclonal anti-Mitochondrial surface p60 (Millipore, MAB1273; IB 1:1000), mouse monoclonal anti-Na^+^/K^+^ ATPase (Santa Cruz, sc-48345; IB 1:1000), mouse monoclonal anti-LAMP1 (H4A3) (Santa Cruz, sc-20011; IB 1:1000; IF 1:400), rabbit polyclonal anti-Calnexin (Santa Cruz, sc-11397; IB 1:1000), mouse monoclonal anti-Galectin 3 (B2C10) (Santa Cruz, sc-32790; IB 1:1000), mouse monoclonal anti-VAPA (4C12) (Santa Cruz, sc-293278; IF 1:400), mouse monoclonal anti-p62/SQSTM (D-3) (Santa Cruz, sc-28359; IB 1:500), mouse monoclonal anti-Alix (BioLegend, 634502; IF 1:200), mouse monoclonal anti-ORP-9 (A7) (Santa Cruz, sc-398961; IF 1:200), HRP-conjugated goat anti-mouse IgG (Thermo Fisher Scientific, 31430; IB 1:5000), HRP-conjugated goat anti-rabbit IgG (Thermo Fisher Scientific; 31460; IB 1:5000), Cyanine Cy™2-conjugated donkey anti-mouse IgG (Jackson ImmunoResearch Laboratories, 715-225-150; IF 1:250), Cyanine Cy™2-conjugated donkey anti-rabbit IgG (Jackson ImmunoResearch Laboratories, 715-225-152; IF 1:250), Cyanine Cy™3-conjugated donkey anti-mouse IgG (Jackson ImmunoResearch Laboratories, 715-162-150; IF 1:250), Cyanine Cy™3-conjugated donkey anti-rabbit IgG (Jackson ImmunoResearch Laboratories, 711-165-152; IF 1:250), Cyanine Cy™5-conjugated donkey anti-mouse IgG (Jackson ImmunoResearch Laboratories, 715-175-150; IF 1:250), and Cyanine Cy™5-conjugated donkey anti-rabbit IgG (Jackson ImmunoResearch Laboratories, 711-175-152; IF 1:250).

### DNA constructs

To create the lentiviral expression construct encoding Halo-tagged EqtSM, the corresponding DNA sequence was PCR amplified from construct pN1-EqtSM_cyto_-Halo^18^ using Phusion high-fidelity DNA polymerase (Thermo Fischer Scientific), inserted via the *EcoRI* and *NotI* sites into dual selection vector pENTR11 (Invitrogen, A10467) and introduced into lentiviral destination vector pLenti CMV hygro DEST (Addgene, 17454) by Gateway cloning according to the manufacturer’s instructions. To create the lentiviral expression construct encoding FLAG-GFP-OSBP, the corresponding DNA sequence was PCR amplified from construct pLJM1-FLAG-GFP-OSBP (Addgene, 134659), inserted via *XmnI* and *EcoRV* sites into pENTR11 and introduced into pLenti CMV hygro DEST by Gateway cloning. To create the expression construct encoding Halo-tagged human calnexin, the ORF of calnexin was PCR amplified from construct mTagBFP2-Calnexin-N-14 (Addgene, 52278) and inserted via *EcoRV* and *EcoRI* sites into expression vector pSEMS-Halo (Covalys Biosciences). The pLenti expression construct encoding Lyso-pHluorin (Addgene, 70113) was described in Rost *et al.*^45^. The expression construct encoding eGFP-tagged P4Mx1 (Addgene, 108121) was described in Zewe *et al.*^50^. The expression constructs encoding eGFP-tagged versions of SPG20 (Addgene, 218578) and SPG20 8xV (Addgene, 218579) were a kind gift from Hemmo Meyer (University of Duisburg-Essen) and described in Ghalot *et al.*^33^. The expression construct encoding mClover-tagged VPS13C (Addgene, 118760) was described in Kumar *et al.*^28^. To create the expression construct encoding mEGFP-tagged VPS13C, the DNA sequence corresponding to mEGFP residues 2-240 was PCR amplified from pSEMS-mEGFP (Covalys Biosciences) and used to exchange the mClover sequence in VPS13C-mClover using Gibson assembly according to the manufacturer’s instructions. To create the expression construct encoding the Halo-tagged ATG2C domain of VPS13C, the DNA sequence corresponding to VPS13C residues 3539 to 3813 was PCR amplified and inserted via the *SfbI* and *XhoI* sites into expression vector pSEMS-Halo (Covalys Biosciences). Single amino acid substitutions were inserted by site-directed mutagenesis (New England Biolabs, E0552S). Respective primers for site-directed mutagenesis, Gibson assembly and cloning are listed in Supplementary Table 1.

### Cell culture and transfection

Human cervical carcinoma HeLa (ATCC CCL-2) and human osteosarcoma U2OS (ATCC HTB-96) cells were cultured in Dulbecco’s modified Eagle’s medium (DMEM) supplemented with 10% FBS (Pan Biotech; P40-47500). Human embryonic kidney 293 cells transformed with Simian Virus 40 large T antigen (HEK293T, ATCC CRL-3216) were cultured in high-glucose Dulbecco’s modified Eagle’s medium (DMEM) containing 2 mM L-glutamine and 10% FBS. Transfections with VPS13C expression constructs were performed using Lipofectamine 3000 (Thermo Fisher, L3000015) according to the manufacturer’s instructions. Transfections with other expression constructs were performed using Linear PEI MAX 40K (Polysciences, Inc., 24765). In brief, 2 µg of plasmid DNA was mixed with 200 µl of 150 mM NaCl and 4 µl of PEI reagent, incubated for 15 min at RT, and then added to cells grown in a 35 mm well. After incubation for 4-5 h at 37°C, cells were washed twice with PBS and cultured overnight in fresh growth medium.

### Lentiviral transduction

For lentiviral transduction of cells with pLenti expression constructs encoding GFP-LAMP1, Lyso-pHluorin, EqtSM-Halo or FLAG-GFP-OSBP, low-passage HEK293T cells were co-transfected with the desired pLenti construct and the packaging vectors psPAX2 (Addgene, 12260) and pMD2.G (Addgene, 12259). After 48 h, the lentivirus-containing medium was harvested, passed through a 0.45 µm filter, mixed 1:1 (v/v) with DMEM containing 8 µg/ml polybrene (Sigma-Aldrich, TR-1003) and used to infect the desired target cell line. At 24 h post-infection, the virus-containing medium was exchanged with DMEM containing the appropriate selection marker (800 µg/ml Hygromycin B, Invitrogen 10687010; 5 μg/ml Blasticidin, Invitrogen 10687010; 1 mg/ml Neomycin, Invitrogen 10687010). After 5 days, positively transduced cells were tested for expression of the desired protein by immunoblot analysis and/or (immuno)fluorescence microscopy.

### Generation of knockout cells

U2OS ATG13-KO cells were a kind gift from Fulvio Reggiori (University of Aarhus, Denmark) and described in Mauthe *et al.*^51^. To knock out VPS13C or PI4K2A in U2OS cells, a mix of CRISPR/Cas9 constructs encoding three different VPS13C or PI4K2A-specific gRNAs and a GFP marker was obtained from Santa Cruz (VPS13C, sc-407531; PI4K2A, sc-429971). The VPS13C-specific gRNA sequences were: A/sense: 5’-TGGCCAGCCTTGACTTTAAA-3’; B/sense: 5’-AGAAGCAGTTGTTGCGACCC-3’; and C/sense: 5’-CAACAACTGCTTCTCCATAA-3’. The PI4K2A-specific gRNA sequences were: A/sense: 5’-CCGCATCGGGCTACCACCAA-3’; B/sense: 5’-CAACATTGTTCCCCGTACAA-3’; and C/sense: 5’-ACTGCTCCAGTTTGAGCGGT-3’. At 24 h post-transfection, single GFP-positive cells were sorted into 96-well plates using a SH800S cell sorter (Sony Biotechnology), expanded, and analyzed for VPS13C and PI4K2A expression by immunoblot analysis.

### Cell lysis and immunoblot analysis

Cells were harvested and lysed in Lysis Buffer (1% TritonX-100, 1 mM EDTA pH 8.0, 150 mM NaCl, 20 mM Tris pH 7.5) supplemented with Protease Inhibitor Cocktail (PIC; 1 μg/ml aprotinin, 1 μg/ml leupeptin, 1 μg/ml pepstatin, 5 μg/ml antipain, 157 μg/ml benzamidine). Nuclei were removed by centrifugation at 600 x g for 10 min at 4°C. Post nuclear supernatants were collected and stored at – 80°C until use. Protein samples were mixed with 2 x Laemmli Sample Buffer (0.3 M Tris HCl, pH 6.8, 10% SDS, 50% glycerol, 10% 2-β-mercaptoethanol, 0.025% bromphenol blue), resolved by SDS-PAGE on 8% (for VPS13C detection) or 12% acrylamide gels and transferred onto nitrocellulose membranes (0.45 μm; GE Health Sciences USA). The membranes were blocked with 5% BSA in PBS for 40 min at RT, washed with 0.05% Tween in PBS (PBST) and then incubated with primary antibody in PBST overnight at 4°C. Next, the membranes were washed three times with PBST and incubated with HRP-conjugated secondary antibody in PBST for 40 min at RT. After washing in PBST, the membranes were developed using enhanced chemiluminescence substrate (ECL; Thermo Fisher Scientific, USA). Images were recorded using a ChemiDoc XRS +System (Bio-Rad, USA) and processed with Image Lab Software (BioRad, USA).

### Lysosome proteomics and lipidomics

#### Affinity purification of lysosomes

A total of 10-12 million HeLa cells stably transduced with LAMP1-GFP were seeded on a 15 cm plate or alternatively on three 10 cm plates in equivalent amounts. Cells were grown in DMEM supplemented with 10% FCS for 3-4 days to a confluency of 70%. For proteomics samples, cells were washed once with PBS and then incubated for 30 min at 37°C in Opti-MEM in the presence or absence of 500 μM LLOMe. For lipidomics samples, cells were washed once with PBS and then incubated for 12 min at 37°C in Opti-MEM in the presence or absence of 1 mM LLOMe. After incubation, the cells were washed twice with ice-cold PBS, detached from the plates by scraping with a rubber policeman in 16 ml ice-cold PBS, transferred to a 50 ml falcon tube on ice and centrifuged at 500 x g for 5 min at 4°C. All steps from here were performed on ice or at 4°C. Cell pellets were gently resuspended in 10 ml ice-cold MIB buffer (10 mM Hepes, 0.21 M mannitol, 0.070 M sucrose, pH 7.5) and centrifuged at 1000 x g for 5 min. This step was repeated once. Next, the cell pellets were resuspended in 0.8 ml freshly prepared and filtered MIB4 buffer (MIB buffer supplemented with 0.5 mM DTT, 0.5% fatty acid-free BSA (Sigma Aldrich), 25 units/ml Benzonase (Sigma Aldrich) and 1 x cOmplete Mini, EDTA-free Protease Inhibitor Cocktail (Roche Diagnostics)), taken up into a 1 ml Luer-Lok™syringe (BD) and passed 25 times through a Balch homogenizer with a tungsten carbide ball of 8.012 mm. The cell lysates were centrifuged at 1,500 x g for 10 min. The supernatants were transferred to fresh tubes and centrifuged again at 1,500 x g for 15 min to generate a post-nuclear supernatant. For affinity purification of lysosomes, anti-GFP IgG microbeads (μMACS™GFP Isolation Kit, Miltenyi Biotec, 130-091-125) were washed in MIB4 buffer and 90 ul of a 1:1 bead:buffer slurry was mixed with 500 ul of postnuclear supernatant. After 1 h of gently mixing on a rotor at an angle of 45°, the suspensions were loaded onto MS columns (Miltenyi Biotec, 130-042-201) that were pre-equilibrated in MIB4 buffer and mounted on a magnetic stand (Miltenyi Biotec, 130-042-109). For the preparation of proteomics samples, the columns were washed once with 500 μl MIB4 buffer and three times with 500 μl MIB buffer. Thereafter, the columns were removed from the magnetic stand and elution was performed with 600 μl MIB buffer. The eluates were divided into 200 μl and 400 μl aliquots for immunoblot and proteomics analysis, respectively. Microbeads were washed once in 200 μl MIB buffer and collected by centrifugation at 21,100 x g for 20 min. For the preparation of lipidomics samples, the columns were washed once with 500 μl MIB4 buffer, three times with 500 μl MIB2 buffer (MIB4 buffer lacking Benzonase and protease inhibitors), and eluted with 600 μl MIB buffer. The eluates were equally divided for immunoblot and proteomics analysis. Microbeads were washed once in 200 μl MIB+ buffer (MIB2 buffer without BSA) and collected by centrifugation at 21,100 x g for 20 min.

#### Immunoblot analysis of purified lysosomes

Semi-quantification of scanned immunoblots of whole cell extracts and purified lysosomes that were loaded on an equivalent basis was performed in Image Lab (version 6.0.1). Using the volume tool, protein bands and corresponding background regions were manually selected for subtraction of background signal. The resulting values for each protein marker were then normalized to those of GFP-LAMP1 from the same sample. For fold-change analysis, control samples were set to 1 and LLOMe-treated sample values were divided by their corresponding controls. Mean values ± SD calculated from four biologically independent experiments were plotted using GraphPad Prism (version 8.0.1).

#### Lysosomal proteomics

Microbead pellets with bound lysosomes were resuspended in 80 μl of 0.5% SDS and incubated at RT for 10 min with gentle shaking. After centrifugation at 21,100 x g for 5 min, the supernatant was transferred to a fresh Eppendorf tube. This was repeated once to avoid any carry over of microbeads. Supernatants were subjected to protein digestion and prepared for proteome analysis using a PreOMICS iST Sample Preparation Kit (Preomics) according to the manufacturer’s instructions. Peptides were resuspended in 10 μl LC-load and subjected to reverse-phase chromatography using a Thermo Ultimate 3000 RSLCnano system connected to a TimsTOF HT mass spectrometer (Bruker Corporation, Bremen) through a captive spray ion source. Peptides were separated on an Aurora Gen3 C18 column (25 cm x 75 μm x 1.6 μm) with CSI emitter (Ionoptics, Australia) at 40°C and eluted using a linear gradient of acetonitrile from 2–35% in 0.1% formic acid at a constant flow rate of 300 nl/min for 44 min followed by an increase in acetonitrile concentration to 50% over 7 min and finally to 85% over 4 min. Eluted peptides were electrosprayed into the mass spectrometer at an electrospray voltage of 1.5 kV and 3 l/min dry gas. MS settings were adjusted to positive ion polarity and an MS range from 100 to 1700 m/z. The scan mode was set to DIA-PASEF. The ion mobility was ramped from 0.7 Vs/cm2 to 1.5 in 100 ms and the accumulation time was 100 ms. 10 MS1 ramps with a window of 26 Da were selected within the mass range 284 to 1385 and the mobility range from 0.7 to 1. 42 [1/k0], resulting in 44 steps per cycle with a mass overlap of 1 Da. Data analysis was performed using MaxQuant (V2.7.0.0 www.maxquant.org)^52,53^ and Perseus (V2.1.5.0 www.maxquant.org/perseus)^54^. Plots were performed with the R software package (www.r-project.org/; RRID:SCR_001905).

#### Lysosomal lipidomics

Microbead pellets with bound lysosomes were subjected to lipid extraction as in Nielsen *et al*.^55^. Briefly, microbeads were resuspended in 200 μl 155 mM ammonium bicarbonate, mixed with 12,5 μl internal lipid standard mix and 988 μl chloroform:methanol 2:1 (v/v), and then shaken in a thermomixer at 2,000 rpm at 4°C for 15 min. After centrifugation (2000 x g, 2 min, 4°C), the lower phase containing lipids was transferred to new tubes and dried in a vacuum centrifuge for 75 min. The dried lipids were resuspended in 100 μl chloroform:methanol 1:2 (v/v) and subjected to shotgun lipidomics as described in Nielsen *et al*.^55^. The samples were analyzed in the negative and positive ionization modes using a Q Exactive Hybrid Quadrupole-Orbitrap mass spectrometer (Thermo Fisher Scientific, Waltham, MA) coupled to TriVersa NanoMate (Advion Biosciences, Ithaca, NY, USA). Data are reported as mol% of total lipids measured.

### Immunofluorescence microscopy

U2OS cells were seeded on glass coverslips at a density of 200,000 cells per 35 mm well in DMEM supplemented with 10% FBS and transfected 24 h after seeding. For experiments with microbeads, the supplier’s stock of 3 μm Polybead® Amino Microspheres was diluted in PBS (1:10, v/v) and resuspended in DMEM containing 10% FBS and 100 U/ml Pen/Strep to 100 µg/ml (for wildtype cells) or 200 µg/ml (for PI4K2A-KO or VPS13C-KO cells) and added to cells at 24h post-transfection. Cells transfected with Halo-tagged constructs were labeled with 50 nM TMR-labeled HaloTag-ligand (Promega, G8252) for 30 min at 37°C and washed five times with culture medium before drug treatment. Cells were treated with 200 nM Apilimod for 2 h and/or 1 mM LLOMe in Opti-MEM for the indicated time. After drug treatment, cells were washed twice with PBS and fixed in 4% (w/v) paraformaldehyde (PFA) for 20 min at RT. After quenching in 50 mM ammonium chloride, cells were permeabilized with permeabilization buffer (PBS containing 0.1% (v/v) Triton X-100 and 1% (w/v) BSA) for 15 min at RT. Immunostaining was performed in permeabilization buffer and nuclei were counterstained with DAPI. Coverslips were mounted onto microscopy slides using ProLong Gold Antifade Reagent (ThermoFisher). Fluorescence images were captured using an Olympus IX-71 DeltaVision Elite microscope equipped with a pco.edge 4.2 (PCO) camera and an Olympus PLAPON 60x (NA 1.42) oil immersion objective. Images were deconvoluted using SoftWoRx software and further processed with Fiji Image J2 software (version 2.3.0/1.53 f; National Institute of Health, USA).

### Live cell imaging

Unless indicated otherwise, live cell imaging was performed using a Zeiss Cell Observer Spinning Disc Confocal Microscope equipped with a TempModule S1 temperature control unit, a Yokogawa Spinning Disc CSU-X1a 5000 Unit, an Evolve EMCCDD camera (Photonics, Tucson), a motorized xyz-stage PZ-2000 XYZ (Applied Scientific Instrumentation) and an Alpha Plan-Apochromat x63 (NA 1.46) oil immersion objective. Filter sets used were: blue emission with BP 445/50, green emission with BP 525/50, orange emission with BP 605/70 and red emission with 690/50. Images were taken using the Zeiss Zen 2012 acquisition software. For transfections with VPS13C expression constructs, U2OS cells were seeded 72 h before imaging at a density of 7,500 cells per well into a µ-Slide 8-well chamber (Ibidi) in DMEM supplemented with 10% FBS. After 24 h, cells were transfected, incubated for 24 h, washed twice with PBS, and incubated with 100 µg/ml pHrodo-microbeads (prepared as described below) in DMEM containing 10% FBS and 100 U/ml Pen/Strep or 1.7 µM 70 kDa dextran in DMEM supplemented with 10% FBS o/n at 37°C. For transfections with other expression constructs, cells were seeded 48 h before imaging at a density of 15,000 cells per well in an µ-Slide 8-well chamber in DMEM supplemented with 10% FBS. After transfection, cells were washed twice with PBS and incubated with 3 µm pHrodo-microbeads or AlexaFluor647-conjugated 70 kDa dextran as inducated above. For cells loaded with dextran, medium was changed to Opti-MEM 2 h prior to imaging after washing with PBS. Halo-tagged constructs were labeled with 30 nM HaloTag® Ligand JaneliaFluor 646 (JF 646) for 15 min in Opti-MEM. Medium was replaced with Opti-MEM supplemented with 30 mM HEPES and 100 U/ml Pen/Strep and then transferred to the stage-top incubator of the microscope preheated to 37°C. The slide was allowed to equilibrate for 10 min and images were taken at every indicated time point with six z-stack of 1 µm. For LLOMe-treatment, medium was exchanged for Opti-MEM containing 1 mM LLOMe, 100 U/ml Pen/Strep and 30 mM HEPES directly at the microscope. For time-lapse recording of hypotonic swelling, cells were imaged in isotonic medium containing 100% Opti-MEM supplemented with 30 mM HEPES and 100 U/ml Pen/Strep. To induce swelling, cells were incubated in 1% Opti-MEM in H_2_O for 5 min and then imaged. Images were processed using Fiji software (NIH, USA).

## Time-lapse recoding of cellular uptake of pHrodo-microbeads

The supplier’s stock of 3 µm Polybead® Amino Microspheres was diluted (1:10, v/v) in 100 mM NaHCO_3_ (pH 8.4) and centrifuged at 10.000 x *g* for 10 min at 10°C. After removal of the supernatant, the microbeads were resuspended in 1 ml of 100 mM NaHCO_3_ (pH 8.4) supplemented with 280 nM pHrodo-Red-succinimidyl ester from a 10 µM stock in DMSO and incubated for 1 h at RT in the dark with continuous shaking. Next, the pHrodo-labeled microbeads were collected by centrifugation at 10,000 x g for 10 min at 10°C, washed four times in 1 ml PBS and stored at 4°C in the dark in the presence of 100 U/ml Pen/Strep. Time-lapse recoding of cellular uptake of pHrodo-microbeads was performed using a Yokogawa CQ1 Confocal Spinning Disk Microscope with temperature, CO_2_, O_2_ and humidity control, a microlense-enhanced dual Nipkow disk system, a camera with a sCMOS chip, a laser-based hardware autofocus and an Olympus IPLSAPO40X2 40x (NA 0.95) air objective. Filter sets used were: green emission with BP 525/50, orange emission with BP 617/73, and red emission with 685/40. Images were taken using the CellPathfinder acquisition software. For the microbead uptake experiment, U2OS cells were seeded at a density of 7,500 cells per well into an µ-Slide 8-well chamber in DMEM supplemented with 10% FBS at 72 h before imaging. After 24 h, cells were transfected with VPS13C-mClover. At 24 h post-transfection, cells were washed once with PBS and incubated in DMEM containing 10% FBS and 100 U/ml Pen/Strep. Next, medium was replaced with Opti-MEM supplemented with 100 U/ml Pen/Strep and cells were transferred to the stage-top incubator of the microscope preheated to 37°C. Right before imaging, 100 µg/ml of pHrodo-microbeads were added in Opti-MEM containing 100 U/ml Pen/Strep. Microbead uptake was monitored for 8 h, with time-lapse images acquired every 1 h. Images were processed using Fiji software (NIH, USA).

### Azido-choline labeling of cells and ER/Golgi-selective SPAAC-reaction

Cells were metabolically labeled with N_3_-Chol using a modified version of the protocol described by Tsuchiya *et al.*^56^. In brief, U2OS or HeLa cells were seeded at a density of 1.5×10^5^ (6-well) or 1.5×10^4^ (8-well) and incubated overnight at 37°C. The next day, the medium was replaced by Opti-MEM containing 500 μM N_3_-Chol (stock 50 mM in H_2_O) and cells were incubated for 24 h to allow metabolic incorporation into endogenous PC. Samples for live-cell imaging and lattice light sheet microscopy were transfected with respective expression constructs prior to incubation with N_3_-Chol. Where indicated, 100 μg/ml of 3 μm pHrodo-microbeads were added to the culture medium. The next day, cells were washed twice with PBS and incubated for 15 min in Opti-MEM containing 100 nM BDP-FL-DBCO (100 μM stock in DMSO) for ER/Golgi-selective labeling of N_3_-Chol-containing PC. Next, cells were washed twice with PBS, incubated in fresh Opti-MEM for 30 min at 37°C and then either harvested for lipid extraction and TLC analysis or imaged by lattice light-sheet microscopy.

### Lipid extraction and TLC analysis

Lipid extraction was performed according to the Bligh-Dyer method^57^. To this end, HeLa cells grown in 6-well dish were trypsinized, resuspended in 1 ml PBS containing 1x protease inhibitor cocktail and centrifuged at 900 x g for 10 min at 4°C. After removal of 900 μl of the supernatant, the cell pellet was resuspended in the remaining 100 μl of PBS, mixed with 375 μl CHCl_3_:MeOH 1:2 (v/v), vortexed and stored at -20°C. Next, the extracts were centrifuged at 21.000 x g for 5 min at 4°C. The supernatant was transferred to a new tube containing 1 vol. (100 μl) CHCl_3_ and 1.25 vol. (125 μl) 0.45 % NaCl (w/v) and vigorously vortexed for 5 min. After centrifugation at 21.000 x g for 5 min at 4°C, the organic lower phase was transferred to a new tube containing 3.75 vol. (375 μl) MeOH:0.45% NaCl and vortexed for 5 min. After centrifugation at 21.000 x g for 5 min at 4°C, the organic lower phase was transferred to a fresh tube and dried to a lipid film in a Christ RVC 2-18 speedvac (Christ, Germany) equipped with a Vacuubrand MZ 2C diaphragm vacuum pump (Vacuubrand, Germany) at 40°C for 40 min. The lipid film was resuspended in 50 μl CHCl_3_:MeOH 1:1 (v/v) and subjected to TLC analysis by spotting 10 μl of the resuspended lipid film on a NANO-ADAMANT HP TLC plate (Macherey & Nagel, Germany) using a Camag ATS4 TLC sampler (CAMAG, Germany). The TLC plate was developed in CHCl_3_:MeOH:NH_3_ 50:25:6 (v/v/v) using a CAMAG ADC2 automatic TLC developer. BDP-FL-DBCO derivatized lipids were visualized with a Typhoon FLA 9500 Biomolecular Imager (GE Healthcare Life Sciences, USA) using a 473 nm excitation laser, LPB filter, 50 μm pixel size and a PMT voltage setting of 290 V. Following fluorescence detection, lipids were stained with iodine vapor to verify that total lipid content between extracts was comparable.

### Lattice light-sheet microscopy

Lattice light-sheet microscopy (LLSM) was performed on a home-built clone of the original design by the Betzig group^58^. Transfected or stably transduced U2OS cells were grown in Opti-MEM containing 500 μM N_3_-Chol and 100 μg/ml 3 μm pHrodo-microbeads on plasma-cleaned and PLL-PEG-RGD-coated^59^ 5 mm round glass coverslips (Art. No. 11888372, ThermoFisher Scientific). After 24 h at 37°C, cells were incubated with 100 nM BDP-FL-DBCO for 15 min at 37°C, washed and incubated in Opti-MEM containing 100 U/ml Pen/Strep for 30 min at 37°C. After that, cells were washed three times with PBS and inserted into a custom-built sample holder, which was mounted onto a piezo stage (sample piezo) for fast sample scan imaging. This ensured that the coverslips were positioned correctly between the excitation and detection objectives inside the sample bath. The latter contained 6 ml of Opti-MEM containing 100 U/ml Pen/Strep at 37°C. Humidity and temperature were precisely controlled by a fully automatic incubation system during the entire experiment (H301-LLSM-SS316 & CO2-O2 Unit BL, Okolab, Italy). For illumination, a dithered square lattice pattern generated by multiple Bessel beams using an inner and outer numerical aperture of the excitation objective of 0.48 and 0.55, respectively, was used. As illumination sources, a 488 nm laser (2RU-VFL-P-300-488-B1R; MPB Communications Inc., Pointe-Claire, Canada) was used for BDP-FL-DBCO, a 561 nm laser (2RU-VFL-P-2000-561-B1R; MPB Communications Inc., Pointe-Claire, Canada) for pHrodo-microbeads, and a 642 nm laser (2RU-VFL-P-2000-642-B1R; MPB Communications Inc., Pointe-Claire, Canada) for HaloTag Ligand JaneliaFluor646 (JF 646)-labeled proteins. The final lattice light-sheet was generated by a water-dipping excitation objective (54-10-7@488-910 nm, NA 0.66, Special Optics), while emitted photons were collected by a water-dipping detection objective (CFI Apo LWD 25XW, NA 1.1, Nikon). Emission was then split via a beamsplitter (FF640-FDi01-25×36, Semrock), filtered by a corresponding dual bandpass filter (Brightline HC 523/610, Semrock) for BDP-FL-DBCO and pHrodo, as well as a longpass filter (647 LP Edge Basic, Semrock) for JF 646. Split emission was detected on individual sCMOS cameras (ORCA-Fusion, Hamamatsu, Japan) with a final pixel size of 103.5 nm. For image acquisition, a sequential two-channel image stack was acquired in sample scan mode by scanning the sample through a fixed light sheet with a step size of 400 nm, which is equivalent to ∼217 nm slicing with respect to the z-axis, considering the sample scan angle of 32.8°. For LLOMe treatment, 1 ml of medium was removed from the sample bath and 1 ml of a freshly prepared 6 mM stock solution of LLOMe in Opti-MEM was added to achieve a final concentration of 1 mM LLOMe. Afterwards, cells were imaged at 4 min intervals.

### Quantification of LLSM data

For quantification of lattice light-sheet microscopy data, raw data were processed using an open-source LLSM post-processing utility called LLSpy (https://github.com/tlambert03/LLSpy) for deskewing, deconvolution, channel registration and transformation. Deconvolution was performed by using experimental point spread functions recorded from 100 nm sized FluoSpheres™ (Art. No. F8803 & F8801 ThermoFisher Scientific) as well as 170 nm PS-Speck™ microspheres (art. No. P7220, ThermoFisher Scientific) and is based on the Richardson–Lucy algorithm using 10 iterations. For channel registration and alignment, 200 nm sized fluorescent TetraSpeck™ microspheres (Art. No. T7280, ThermoFisher Scientific) were imaged with a step size of 350 nm. Alignment was done with a 2-step transformation method that deploys an affine transformation in xy and a rigid transformation in z.

For line profiles of internalized pHrodo-microbeads in Fiji, z-stacks at each time point were selected so that the center cross-section of the microbeads was in the focal plane. Then, an arrow was drawn at the indicated position using the arrow tool with a width of “5” in Fiji. Intensity profiles for each channel (EqtSM-Halo, pHrodo, cPC_ER_) and time point were exported to Excel, and the maximum value of each channel at time point 0 min was set to 100. Every following time point was normalized to this baseline. Graphs displaying the normalized data were generated using GraphPad Prism software version 8.3.0 (GraphPad Software LLC, USA).

For quantification of the signal development of CNX-Halo and cPC_ER_ around pHrodo-microbead-containing lysosomes over time, z-stacks at each time point were selected so that the center cross-section of the microbeads was in focus. Then, a region of interest (ROI) was manually drawn around each bead for every time point in Fiji. Everything outside this region was removed using the “Clear outside” function to prevent possible artifacts in the quantification. Then, a modified version of the ImageJ macro described in Maib and Murray^60^ was used to measure intensities in CNX-Halo and cPC_ER_ channels, using the pHrodo signal as source channel for creating a mask. The modified macro was generated by Steffen Wolke-Hanenkamp (Osnabrück University) and is provided in the Supplementary Information. If required, the signal in the pHrodo channel was adjusted, ensuring detection by the macro. CNX-Halo and cPC_ER_ channels in each individual bead were normalized by setting the time point 0 min to 100% and each subsequent time point was normalized by calculating the relative change compared to the 0 min timepoint. To assess relative changes between the two channels, a ratio was calculated by dividing the normalized intensity of cPC_ER_ by the normalized intensity of CNX-Halo, performed individually for each timepoint for each bead. Statistical significance was assessed using an unpaired two-tailed t-test using Welch’s correction to account for unequal variances using GraphPad Prism software version 8.3.0 (GraphPad Software LLC, USA).

Quantification of cPC_ER_ and EqtSM-Halo accumulation on pHrodo-bead containing lysosomes was performed using Imaris 9.5 software (Bitplane). For this analysis, individual cells were isolated and cropped from rotated and deconvoluted LLSM hyperstack images in ImageJ while ensuring that internalized pHrodo-beads were present in all acquired timepoints. These hyperstack image files were then loaded into Imaris 9.5 using the corresponding ImageJ plugin option to create a 3D reconstruction of each cell. Using the pHrodo channel as a source, a surface was rendered with a surface detail of 0.5 μm. The threshold absolute intensity was manually adjusted to cover the entire bead structure across all time points. The generated surfaces were then filtered by the number of voxels to exclude any surfaces not representing pHrodo-microbeads. Statistics showing the mean intensity at the surface of all channels (EqtSM-Halo, pHrodo, cPC_ER_) at each time point were exported into Excel for analysis. For each channel and time point, all microbeads within a cell were treated as a single surface and averaged. Timepoint 0 min was used to normalize the subsequent time points. Graphs were created using GraphPad Prism software.

### Time-lapse recording of lysosomal repair

Time-lapse recordings of cells exposed to lysosome-damaging drugs were performed using a Zeiss Cell Observer Spinning Disc Confocal Microscope equipped with a TempModule S1 temperature control unit, a Yokogawa Spinning Disc CSU-X1a 5000 Unit, an ORCA Flash 4.0 V3 camera (Hamamatsu), a motorized xyz-stage PZ-2000 XYZ (Applied Scientific Instrumentation) and an Alpha Plan-Apochromat x63 (NA 1.46) oil immersion objective. The following filter combinations were used: blue emission with BP 445/50, green emission with BP 525/50, orange emission BP 605/70. All images were acquired using Zeiss Zen 2012 acquisition software. At 24 h before imaging, U2OS wildtype and VPS13C-KO cells stably transduced with LysopHluorin were seeded into a µ-Slide eight-well glass-bottom chamber (Ibidi, 80827) at a density of 16.000 cells per well in DMEM supplemented with 10% FBS. After 24 h, medium was replaced with Opti-MEM containing 30 mM HEPES and 100 U/ml Pen/Strep. At 10 min before imaging, medium was replaced with imaging medium (IM) containing 30 mM HEPES, 140 mM NaCl, 2.5 mM KCl, 1 mM MgCl_2_, 1.8 mM CaCl_2_ and 10 mM D-glucose, pH 7.4. Cells were immediately transferred to a stage-top incubator preheated to 37°C. The slide was allowed to equilibrate for 10 min before initiation of image acquisition. After positions were obtained, cells were treated with 1mM LLOMe for 10 min, washed once with IM, and chased for 2 h in fresh IM. Images were captured immediately before treatment, at 10 min of LLOMe treatment, and right after the 2 h chase. For experiments with LysoTracker-labeled cells, medium was exchanged for IM containing 75 nM LysoTracker (LT) and cells were immediately transferred to a stage-top incubator preheated to 37°C. The slide was allowed to equilibrate for 10 min before initiation of image acquisition with the Zeiss Cell Observer SD microscope. Time-lapse images were acquired every 60 s (six z-sections, 1 μm apart). After 3 min of image acquisition, GPN was directly added into the well to a final concentration of 200 μM without pausing image acquisition. After 2 min of GPN exposure, acquisition was paused for 2 min to aspirate the GPN-containing medium, the cells were washed once with LT containing IM and fresh LT-containing IM was added before acquisition was resumed.

### Quantitative assessment of lysosomal repair

To quantitatively assess lysosomal repair, the number of LysopHluorin-positive puncta in cells was determined after setting a manual threshold to subtract background fluorescence. Puncta within proximity were separated using the watershed function. Next, for each time point, all puncta with pre-determined characteristics were counted automatically (size 0.1–5 µm^2^, circularity 0.5-1). The number of LysoTracker-positive puncta were counted after setting an automatic threshold at t = 0 min to subtract background fluorescence. Puncta within proximity were separated using the watershed function. Next, for each time point, all puncta with pre-determined characteristics were counted automatically (size 0.2–5 µm^2^, circularity 0.5-1). For normalization of the number of puncta per cell area, the number of obtained puncta for each time point was divided by the total measured area to account for differences in cell size and then multiplied by 100 to obtain the number of puncta per 100 µm^2^ cell area. For normalization relative to the maximal value, the number of puncta per cell per time point was determined and divided by the maximum value. Image J macros used for the quantification of LysopHluorin and LysoTracker puncta are provided in Supplementary Information.

### Statistics and reproducibility

Each experiment was repeated at least twice with similar results using independent experimental samples and statistical tests as specified in the figure legends. Source data with sample sizes, number of technical and/or biological replicates, means, standard deviations, and calculated p values (where applicable) are provided in the Source Data file.

## Data availability

All data generated or analysed during this study are included in the manuscript and supporting file. Source Data files have been provided for Figures 1-8 and Supplemental Figures 1, 7 and 8. The mass spectrometry proteomics data have been deposited to the ProteomeXchange Consortium via the PRIDE partner repository with the dataset identifier PXD069825^61,62^.

## Acknowledgements

We gratefully acknowledge Sergei M. Korneev (Osnabrück University) for synthesizing azido-choline, Murali Amalai (Osnabrück University) for the energy minimization model of DBCO-modified PC, Steffen Wolke-Hanenkamp (Osnabrück University) for creating a modified Fiji/ImageJ macro for image analysis, Stefan Walter (Osnabrück University) for technical assistance with LC-MS/MS, Fulvio Reggiori (Aarhus University) for providing U2OS ATG13-KO cells, and Hemmo Meyer (University of Duisburg-Essen) for providing DNA constructs.

## Author contributions

J. C. M. H. designed the research and wrote the manuscript with critical input from O. W. A., C. S., E. S. and F. B.; O. W. A., C. S., E. S. and F. B. performed the experiments and analyzed the results with critical input from K. S., E. D., A. K. L. and L. S. P.; B. F. created DNA constructs; A. H. established the majority of transduced cell lines; R. K. and M. H. assisted C. S. and E. D. with lattice light-sheet microscopy experiments and image processing; K. S. assisted F. B. with proteomics experiments with critical input from F. F. and B. M. E.; O. W. A. and K. M. performed the lipidomics experiments and analyzed the results; all authors critically discussed results and commented on the manuscript.

## Code availability statement

All custom code is available in the Online Methods section.

