## Supplemental Information for "VPS13C/PARK23 initiates lipid transfer and membrane remodeling for efficient lysosomal repair"

**This PDF includes:**

Supplementary Figure 1 - 9
Supplementary Table 1
Image J Macros
Uncropped blots

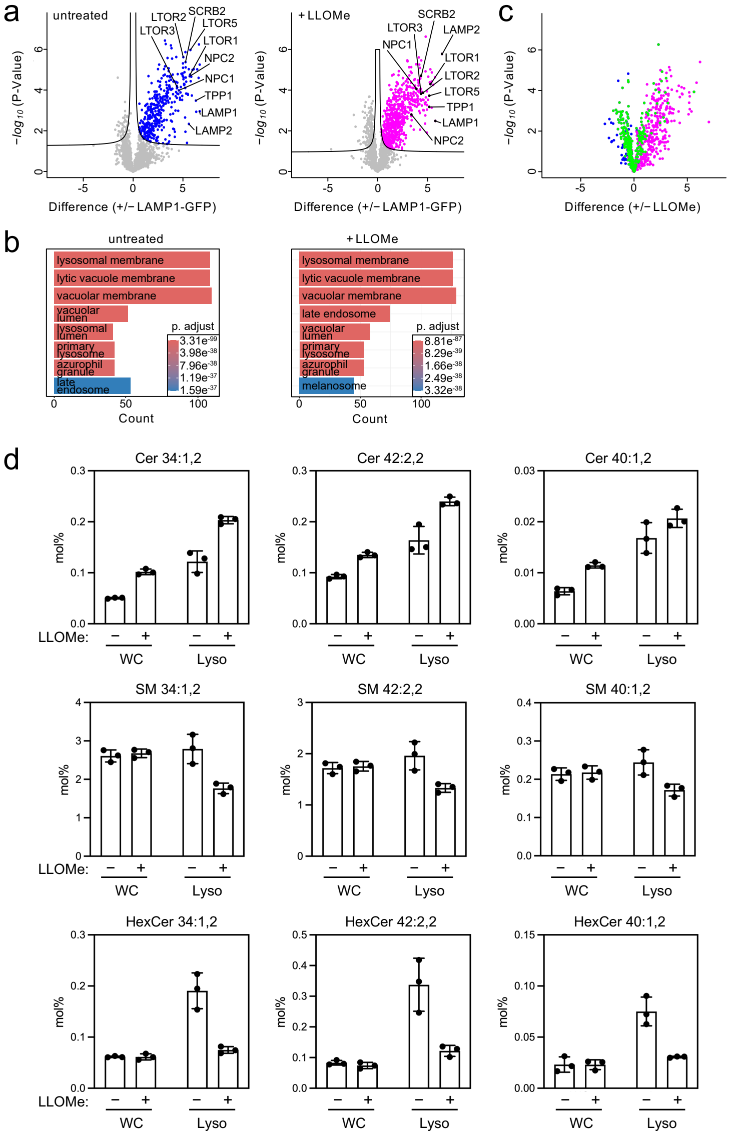


**Supplementary Figure 1. Protein and lipid composition of lysosomes affinity-purified from control and LLOMe-treated HeLa cells.**

(**a**) Volcano plots showing enrichment of various lysosomal membrane proteins in lysosome isolates from untreated or LLOMe-treated (500 µM, 30 min) LAMP1-GFP-expressing HeLa cells relative to background HeLa cells. Fold changes were calculated from three independent biological replicates and plotted on the x-axis against the negative logarithmic *P*-values on the y-axis.

(**b**) Gene Ontology (GO) enrichment analysis of lysosome isolates relative whole cells. Shown are data from LAMP1-GFP-expressing HeLa cells treated as in (a).

(**c**) Volcano plot of proteins enriched in lysosome isolates from untreated and LLOMe-treated LAMP1-GFP HeLa cells and grouped according to how lysosomal damage affects their relative levels. *Green*, proteins enriched in both intact (–LLOMe) and damaged (+LLOMe) lysosomes; *blue*, proteins selectively depleted in damaged lysosomes; *magenta*, proteins selectively enriched in damaged lysosomes.

(**d**) Lipid composition of whole cell lysates (WC) and lysosomes purified from untreated and LLOMe-treated (500 µM, 30 min) HeLa cells expressing LAMP1-GFP was determined by mass spectrometry-based shotgun lipidomics. Levels of the different lipid species are expressed as mol% of total identified lipids. SM, sphingomyelin; Cer, ceramide; HexCer, hexosylceramide.

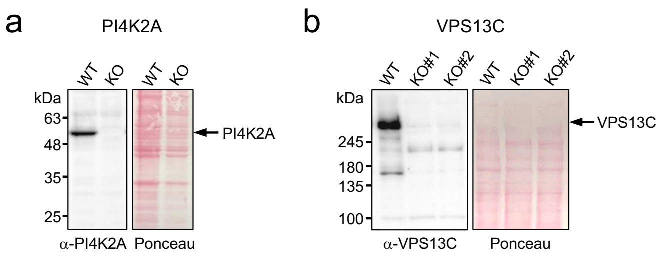


**Supplementary Figure 2. Validation of PI4K2A-KO and VPS13C-KO cell lines by immunoblot analysis.**

(**a**) U2OS cells lacking PI4K2A were created by CRISPR/Cas9. Loss of PI4K2A was confirmed by immunoblot analysis with an anti-PI4K2A antibody, using Ponceau S staining as loading control. Migration of the PI4K2A protein is marked by an arrow.

(**b**) U2OS cells lacking VPS13C were created by CRISPR/Cas9. Loss of VPS13C was confirmed by immunoblot analysis with an anti-VPS13C antibody, using Ponceau S staining as loading control. Migration of the VPS13C protein is marked by an arrow.

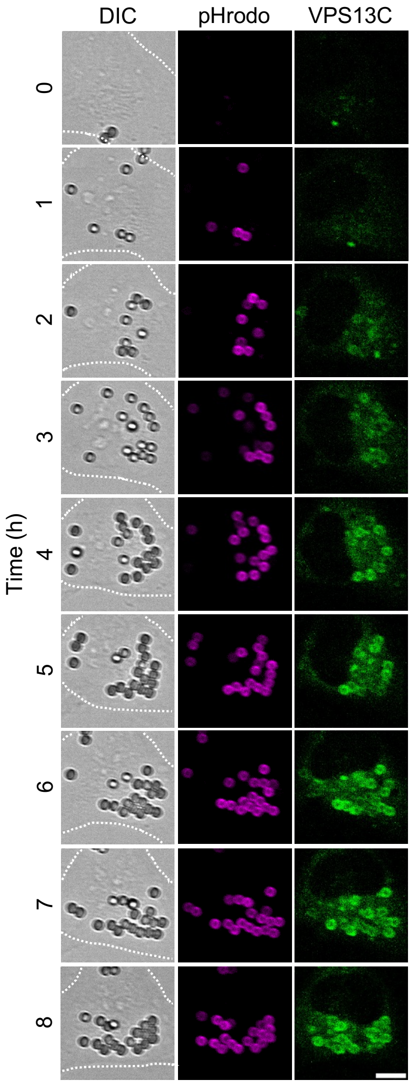


**Supplementary Figure 3. pHrodo-conjugated beads internalized by U2OS cells readily gain pHrodo fluorescence and induce large-scale mobilization of VPS13C.**
Time-lapse images of U2OS cells transfected with VPS13C-mClover (*green*) and incubated with 3 µm pHrodo-labeled beads (*magenta*). Cells were imaged by spinning disk microscopy. Scale bar, 10 µm.

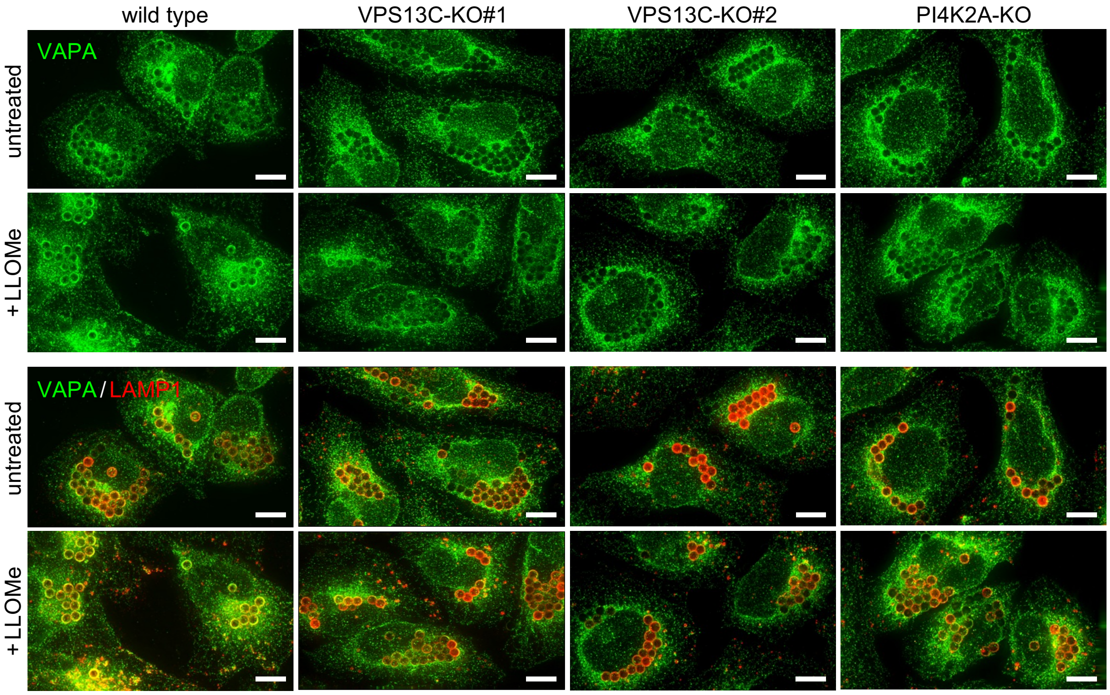


**Supplementary Figure 4. VPS13C is required for tethering damaged microbead-containing lysosomes to the ER.**
U2OS wildtype (WT), VPS13C-KO1, VPS13C-KO2 and PI4K2A-KO cells were fed 3 μm polystyrene beads, treated with LLOMe (1 mM, 20 min) as indicated, immunostained for LAMP1 (*red*) and VAPA (*green*), and imaged by DeltaVision microscopy. Scale bar, 10 μm.

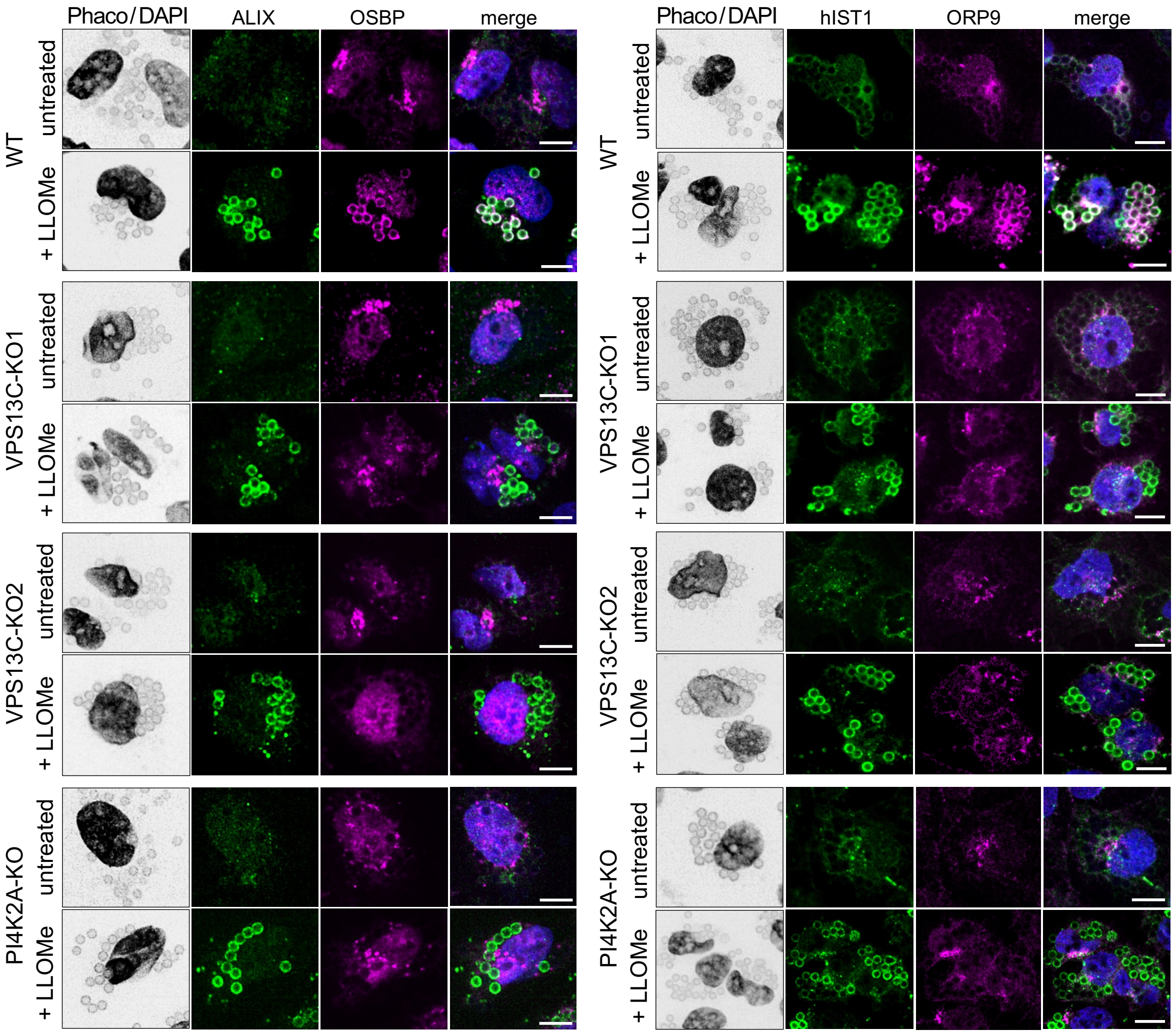


**Supplementary Figure 5. VPS13C is essential for recruiting OSBP and ORP9 to damaged lysosomes.**

Phase contrast (Phaco) and fluorescence images of polystyrene bead-containing untreated or LLOMe-treated (1 mM, 10 min) U2OS wildtype (WT), VPS13C-KO1, VPS13C-KO2 or PI4K2A-KO cells immunostained for OSBP (*magenta*) and ALIX (*green*) or ORP9 (*magenta*) and hIST1 (*green*) and counterstained with DAPI (*blue*). Cells were imaged by spinning disk microscopy. Scale bar, 10 µm.

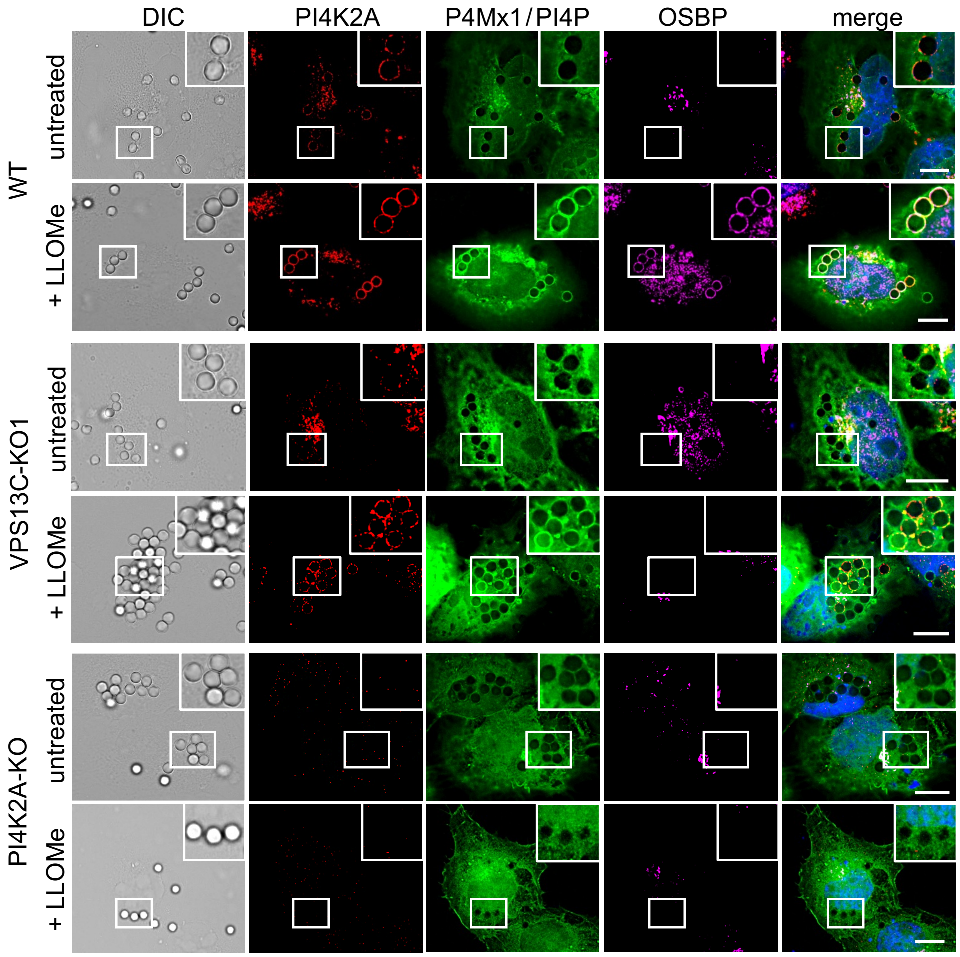


**Supplementary Figure 6. Mobilization of PI4K2A to damaged lysosomes is independent VPS13C.** Differential interference contrast (DIC) and fluorescence images of polystyrene bead-containing untreated or LLOMe-treated (1 mM, 10 min) U2OS wildtype (WT), VPS13C-KO1 or PI4K2A-KO cells expressing P4MX1-GFP (*green*) and immunostained for PI4K2A (*red*) and OSBP (*magenta*) and counterstained with DAPI (*blue*). Cells were imaged by DeltaVision microscopy. Scale bar, 10 µm.

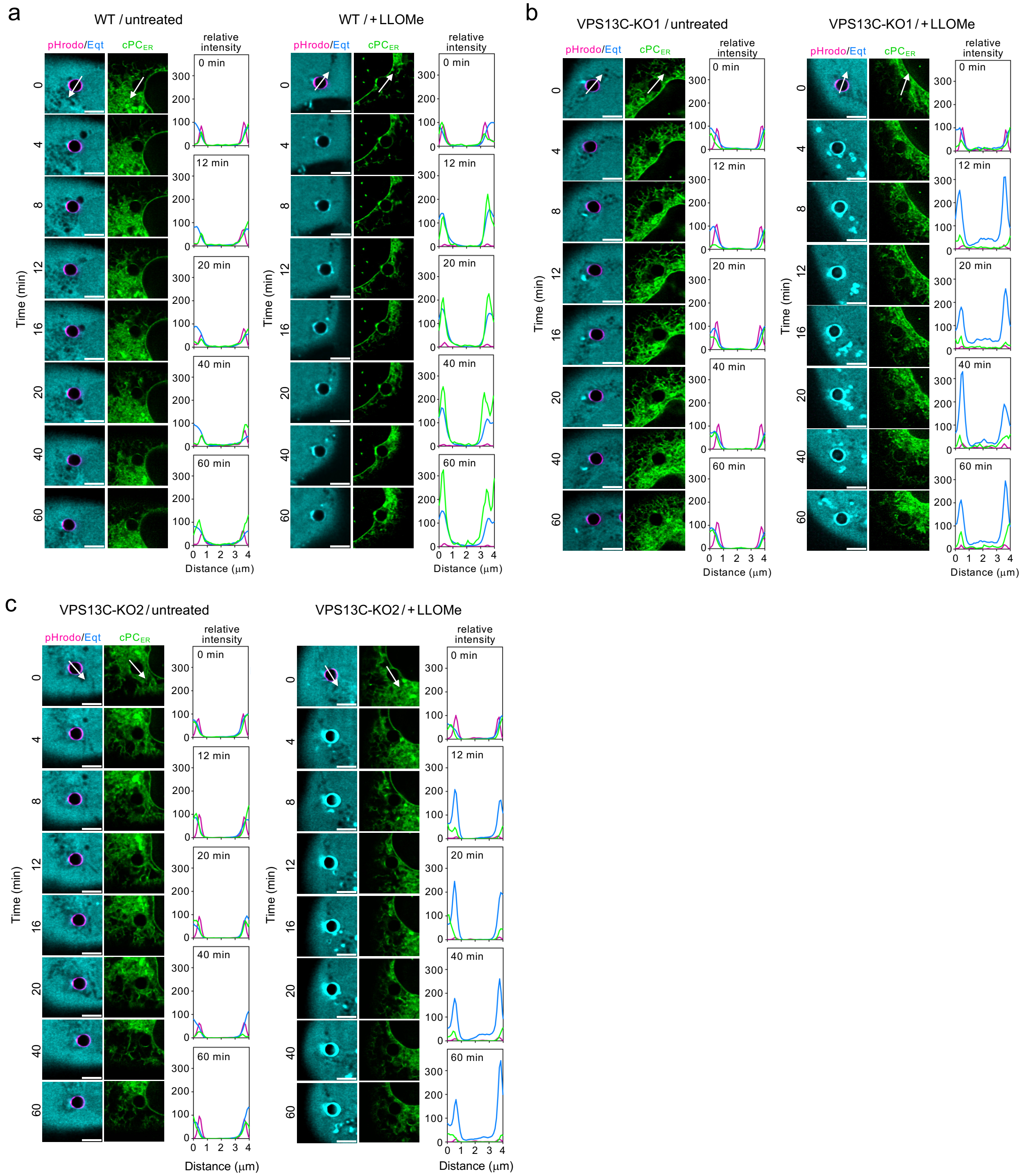


**Supplementary Figure 7. VPS13C removal disrupts delivery of cPC$_{ER}$ to damaged lysosomes.** (**a, b, c**) U2OS wildtype, VPS13C-KO1 and VPS13C-KO2 cells expressing EqtSM-Halo (*cyan*) were fed pHrodo-beads (*magenta*), incubated with $N_3$-Chol, stained with ER-DBCO (cPC$_{ER}$, *green*) and then subjected to time-lapse imaging in the absence (untreated) or presence of 1 mM LLOMe. Cells were imaged by LLSM and only zoom-ins of imaged cells are shown. Line scans show the intensity profiles of EqtSM-Halo (*cyan*), pHrodo (*magenta*) and cPC$_{ER}$ (*green*) signals along the path of the arrows. Profiles are plotted as relative intensities for each channel normalized to the 0 min time point.

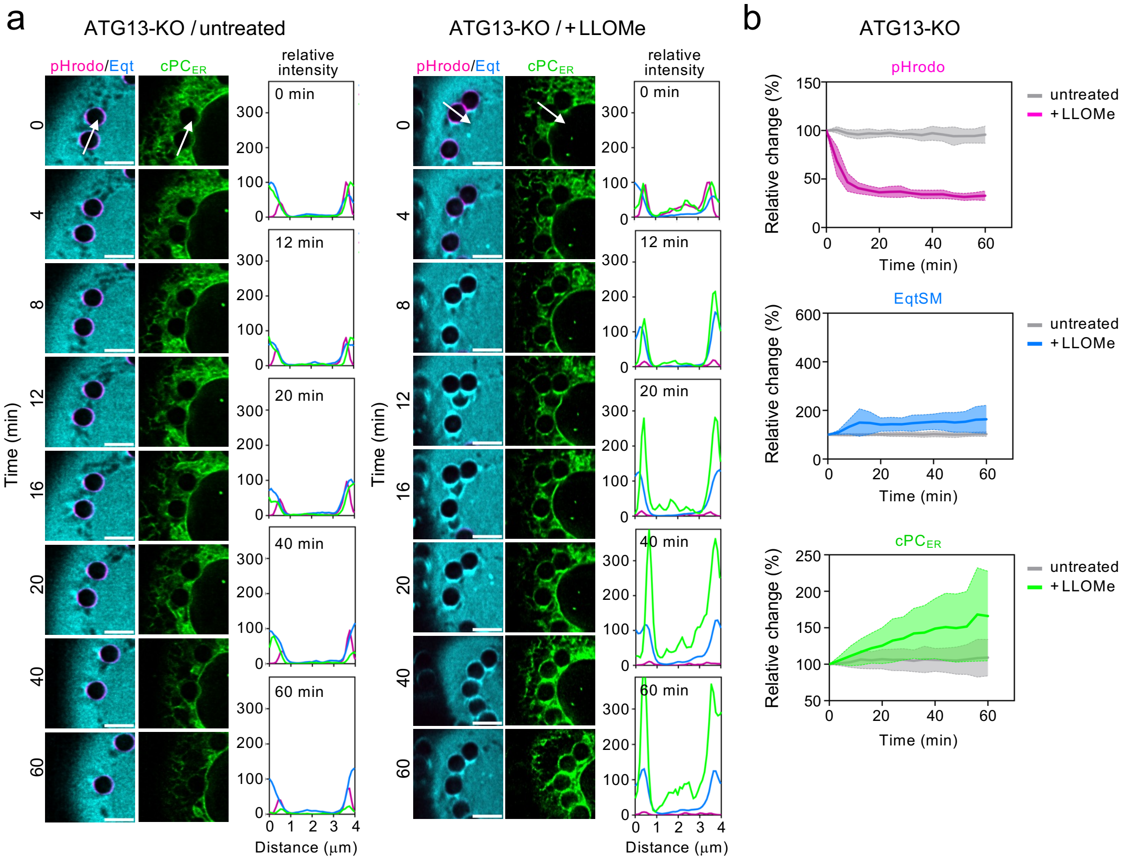


**Supplementary Figure 8. Accumulation of cPC$_{ER}$ on damaged lysosomes is not affected in cells defective in autophagy.**

(**a**) U2OS ATG13-KO cells expressing EqtSM-Halo (*cyan*) were fed pHrodo-beads (*magenta*), incubated with $N_3$-Chol, stained with ER-DBCO (cPC$_{ER}$, *green*) and then subjected to time-lapse imaging in the absence (untreated) or presence of 1 mM LLOMe. Cells were imaged by LLSM and only zoom-ins of imaged cells are shown. Line scans show the intensity profiles of EqtSM-Halo (*cyan*), pHrodo (*magenta*) and cPC$_{ER}$ (*green*) signals along the path of the arrows. Profiles are plotted as relative intensities for each channel normalized to the 0 min time point. Scale bar, 5 µm.

(**b**) Time course plotting the relative changes in pHrodo, EqtSM-Halo, and cPC$_{ER}$ signals on the surface of pHrodo-bead containing lysosomes in U2OS ATG13-KO cells treated as in (a). A 3D-surface was generated around the pHrodo-positive beads using Imaris software, and the mean fluorescence intensities of the respective channels on this surface were quantified. For each cell, signals from all bead-associated surfaces were averaged. Values were normalized to the signal at timepoint 0 min and plotted as mean ± SD over time. ATG13-KO (*n* = 3): control, 10 cells; +LLOMe, 9 cells.

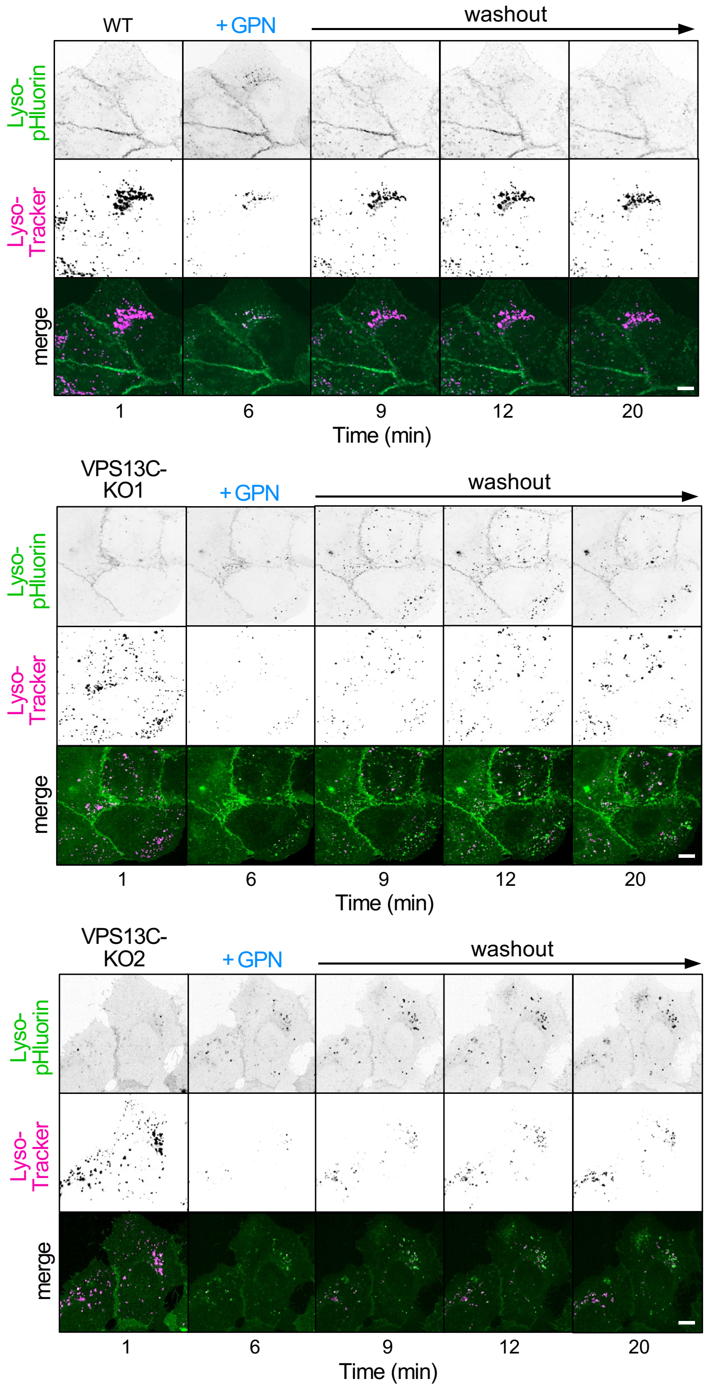


**Supplementary Figure 9. VPS13C removal disrupts lysosomal repair.**
Time-lapse images of LysoTracker-labelled (*magenta*) and Lyso-pHluorin-expressing (*green*) U2OS wildtype (WT), VPS13C-KO1 and VPS13C-KO2 cells pulse-treated with GPN (200 μM, 2 min). Cells were imaged by spinning disk microscopy. Scale bar, 10 μm.

**Supplementary Table 1. Primers used for cloning and site-directed mutagenesis.**

| Primer name | Primer sequence (5'-3') |
|---|---|
| hCalnexin-ORF_fwd | gaatgatatcatggaagggaagtggttgctgtgtatg |
| hCalnexin-ORF_rev | gttagaattcctctcttcgtggctttctgtttcttgg |
| EqtSM$_{cyto}$-Halo_fwd | cacagaattcaccatgtccgcagacgtgg |
| EqtSM$_{cyto}$-Halo_rev | cacacagcggccgcctaaccggaaatctccagagtagac |
| VPS13C ORF_fwd | gctgtacaaggaccatctcaaagaacaagaag |
| VPS13C ORF_rev | tgctcacgccaggtacacttgaaacatcaac |
| Halo-ATG2C_SbfI_fwd | cacacctgcaggtgaattaatggagacttcaatgactg |
| Halo-ATG2C_XhoI_rev | cacactcgagcctatgccctctgaatgccttg |
| VPS13C-V3563Q_fwd | tgtgggtgctcaggcccgtc |
| VPS13C-V3563Q_rev | agcccttttccaattcctttaaagaatc |
| FLAG-GFP-OSBP_fwd | cacagaaccaattcctaccggtgccatggactac |
| FLAG-GFP-OSBP_rev | cacagatatcgtcgagaattgatccgcgc |
| mEGFP w/o Met_fwd | aagtgtacctggcgtgagcaagggctgagcaagggcg |
| mEGFP w/o Met_rev | tgagatggtccttgtacagctcgtccatgc |

#### Image J Macros

Quantification of VPS13C-mClover, GFP-OSBP, and LysopHluorin puncta

1. //set Threshold manually
2. setOption("BlackBackground", false);
3. run("Convert to Mask", "method=Default background=Dark");
4. run("Fill Holes","stack");
5. run("Watershed", "stack");
6. run("Analyze Particles...", "size=0.1-5 circularity=0.50-1.00 show=Outlines display exclude summarize stack");
7. run("Next Slice [>]")
8. //Repeat step 6 and 7 until all time points are analyzed

Quantification of LysoTracker puncta

1. run("Subtract...", "value=20 stack");
2. setAutoThreshold("Default dark no-reset");
3. setOption("BlackBackground", false);
4. run("Convert to Mask", "method=Default background=Dark");
5. run("Convert to Mask", "method=Default background=Light");
6. run("Watershed", "stack");
7. run("Analyze Particles...", "size=0.2-5 circularity=0.5-1.00 show=Outlines display exclude summarize stack");
8. run("Next Slice [>]")
9. // repeat step 7 and 8 until all time frames have been quantified

Quantification of pHrodo, cPC$_{ER}$ andCNX-Halo and  EqtSM-Halo signals on pHrodo-microbead containing lysosomes

Macro based on Maib, H. and Murray, D.H. (2022) ‘A mechanism for exocyst-mediated tethering via Arf6 and PIP5K1C-driven phosphoinositide conversion’, *Current Biology*, 32(13), pp. 2821-2833.e6 (https://doi.org/10.1016/j.cub.2022.04.089) with modifications by Steffen Wolke-Hanenkamp (Ultraphysics Division, Osnabrück University).

```
// Check for the Excel plugin
if (File.exists(getDirectory("plugins") + "/Read_and_Write_Excel-1.1.7.jar") != 1) {
   exit("ResultsToExcel Plugin not found. Make sure you activated the plugin in the
ImageJ updater. ('Help' > 'Update...' > 'Manage update sites')");
}
// Get input image and basic parameters
dir1 = getDirectory("image");
name = getTitle();
selectWindow(name);
getDimensions(width, height, channels, slices, frames);
run("Split Channels");
// Open results Excel sheet
run("Read and Write Excel", "file_mode=read_and_open file=[" + dir1 +
"/CS_BeadsResults.xlsx] sheet=[C1]");
// Iterate over all time frames
for (i = 1; i <= frames; i++) {
   // --- MASK GENERATION (Channel 2) ---
   selectWindow("C2-" + name);
   Stack.setFrame(i);
   run("Duplicate...", "title=C2-mask-frame_" + i + ".tif frames=" + i);
   maskID = getImageID();
   setAutoThreshold("Li dark");
   run("Convert to Mask", "method=Li background=Dark calculate");
   setOption("BlackBackground", false);
   run("Fill Holes");
   run("Watershed");
   run("Analyze Particles...", "size=5-250 circularity=0.1-1.00 show=Overlay clear add
stack");
   // --- ROI ENLARGEMENT ---
   upperROI = roiManager("count");
   for (index = 0; index < upperROI; index++) {
      roiManager("Select", index);
      run("Enlarge...", "enlarge=-1");
      run("Make Band...", "band=1.25");
      roiManager("Update");
   }
   // --- MEASURE CHANNEL 1 ---
   selectWindow("C1-" + name);
```

```
    Stack.setFrame(i);
    run("Duplicate...", "title=C1-double.tif frames=" + i);
    C1doubleID = getImageID();
    roiManager("Deselect");
    roiManager("Measure");
    run("Read and Write Excel", "file_mode=queue_write dataset_label=[Frame " + i + "]
sheet=[C1]");
    run("Clear Results");
    close(C1doubleID);
    // --- MEASURE CHANNEL 2 ---
    selectWindow("C2-" + name);
    Stack.setFrame(i);
    run("Duplicate...", "title=C2-double.tif frames=" + i);
    C2doubleID = getImageID();
    roiManager("Deselect");
    roiManager("Measure");
    run("Read and Write Excel", "file_mode=queue_write dataset_label=[Frame " + i + "]
sheet=[C2]");
    run("Clear Results");
    close(C2doubleID);
    // --- MEASURE CHANNEL 3 ---
    selectWindow("C3-" + name);
    Stack.setFrame(i);
    run("Duplicate...", "title=C3-double.tif frames=" + i);
    C3doubleID = getImageID();
    roiManager("Deselect");
    roiManager("Measure");
    run("Read and Write Excel", "file_mode=queue_write dataset_label=[Frame " + i + "]
sheet=[C3]");
    run("Clear Results");
    close(C3doubleID);
    // --- SAVE MASK ---
    selectImage(maskID);
    saveAs("PNG", dir1 + File.separator + i + "C2-Masks");
    close(maskID);
}
// Finalize Excel export and clean up
run("Read and Write Excel", "file_mode=write_and_close");
close("*");
close("Results");
close("ROI Manager");
// Notify user
waitForUser("End", "Macro has finished");
```

### Uncropped blots

#### Figure 1c

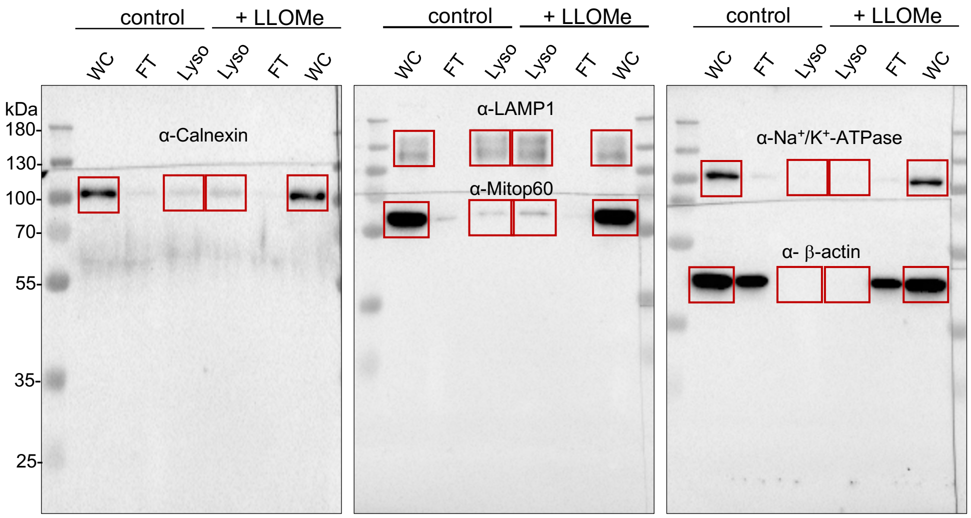


#### Figure 2b

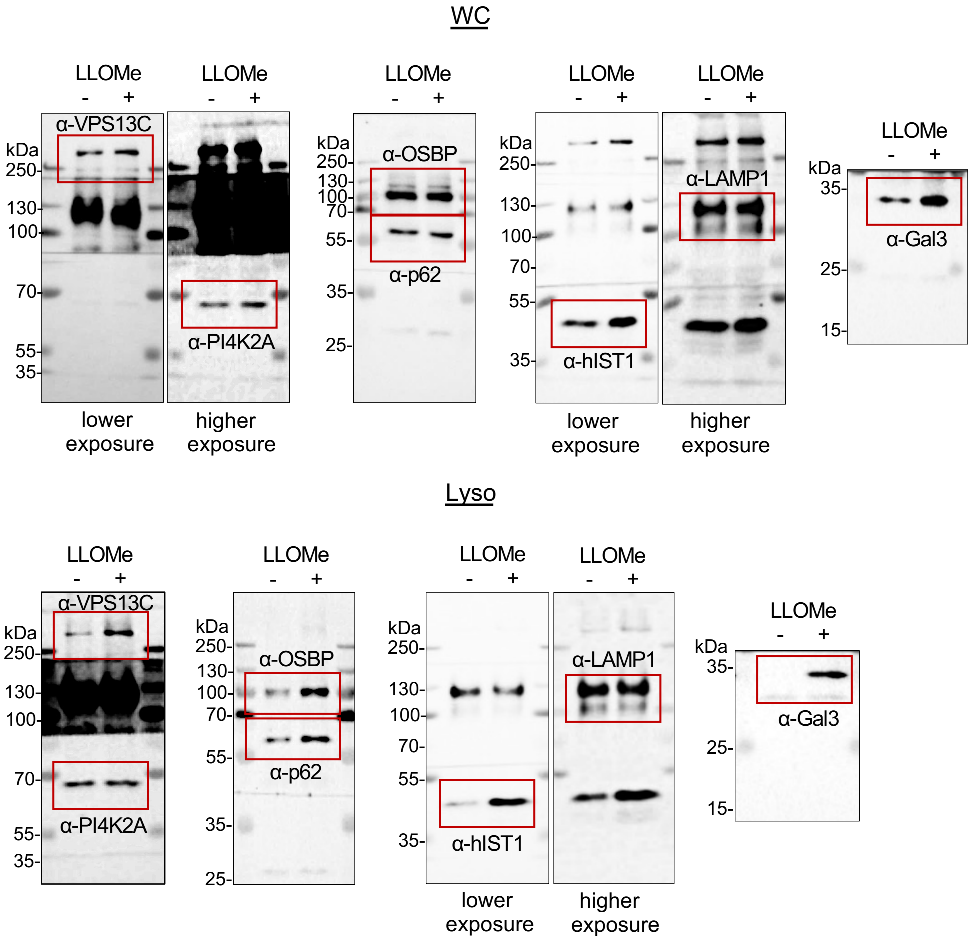
